# Hyperbolic disc embedding of functional human brain connectomes using resting state fMRI

**DOI:** 10.1101/2021.03.25.436730

**Authors:** Wonseok Whi, Seunggyun Ha, Hyejin Kang, Dong Soo Lee

## Abstract

The brain presents a real complex network of modular, small-world, and hierarchical nature, which are features of non-Euclidean geometry. Using resting-state functional magnetic resonance imaging (rs-fMRI), we constructed a scale-free binary graph for each subject, using internodal time-series correlation of regions-of-interest (ROIs) as a proximity measure. The resulted network could be embedded onto manifolds of various curvature and dimensions. While maintaining the fidelity of embedding (low distortion, high mean average precision), functional brain networks were found to be best represented in the hyperbolic disc. Using 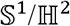 model, we reduced the dimension of the network into 2-D hyperbolic space and were able to efficiently visualize the internodal connections of the brain, preserving proximity as distances and angles on the hyperbolic discs. Each individual disc revealed decentralized nature of information flow and anatomic relevance. Using the hyperbolic distance on the 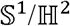 model, we could detect the anomaly of network in autistic spectrum disorder (ASD) subjects. This procedure of embedding grants us a reliable new framework for studying functional brain networks and the possibility of detecting anomalies of the network in the hyperbolic disc on an individual scale.

## Introduction

Significant progress has been achieved over the last decades in unveiling the structure and mechanism of the human brain, which is one of the most complicated systems of nature. Especially, studies on the network geometry, which offer a new perspective on the framework of structural brain networks^1,2^, or connectomes, have revealed that structural brain networks share certain topological properties such as modularity^3^, small-world^4^, and heavy-tailed degree distribution^5^.

However, Euclidean geometry, which serves as a standard framework of investigation in our physical reality, does not seem to suffice in explaining the observed connectivity between the brain regions, functional as well as structural. Instead, an increasing amount of evidence suggests that the geometry of negative curvature i.e. hyperbolic geometry serves as the more appropriate explanatory platform of the above-mentioned topologic features of the structural brain networks that dominates the formation of the network and the probability of connections between the brain regions^6,7^.

Since the functional brain network was closely related to the connectomes^8–10^, we assume that the functional brain network best fits with the hyperbolic geometry, which is regarded as the geometric representation of high-dimensional tree or tree-like graphs with hierarchies^11^. However, real complex graphs could also have a cycle or torus-like structures therein in any dimensions and necessitated the trial-fitting to spherical space of positive curvatures. As hyperbolic and spherical spaces with various curvatures and Euclidean space can be combined to represent real complex graphs with the product of spaces, any real complex graphs can now be tested for their feasibility of fitting to the model of a product of spaces with various negative and/or positive curvatures. The only reserve is to find the model space of appropriate dimension and curvature to fit the real complex networks.

By this means, we later show that we successfully embedded the functional brain graphs observed on resting-state functional MRI (rs-fMRI) to be fit to the 2-dimensional hyperbolic plane (disc) with better fidelity compared with other product manifolds of low dimension. In this study, we used a purely geometric 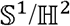 model, which is well known to reveal the hidden geometry of many other real complex networks of non-Euclidean nature, such as internet connection^12^, world trade web^13^ or even the structural brain network^6,7^.

When we adopted the binary graph obtained by rs-fMRI as an input to the 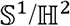 model, we could fit the nodes to their optimal positions on the hyperbolic disc proposed by Papadopoulos et al.^14^ and further publicized as *Mercator*^15^. The position of the nodes could be used to calculate the hyperbolic distance between each node, which serves as a unique variable for determining the connection probability between the nodes on this disc^16^. We used the distance matrix for comparison of the proximity between brain region pairs across the individuals with autism spectrum disorder (ASD) and control group to see if the method is applicable to real cases of diseased subjects.

## Results

### Distribution of time-series correlation in rs-fMRI

We used the pre-defined parcellation of the whole brain into 274 regions of interest (ROIs) and computed the time-series correlation between the ROIs. The distribution of the value of coefficients is shown in (**Fig. 1a**). Overall, a positive correlation was dominant among edges from the healthy young adults, while a small portion of edges showed a negative correlation. Detailed visualization of correlation matrix from a representative case is shown in (**Fig. 1b**). We regarded that a certain degree of anticorrelation between brain regions also suggested connection and took the absolute value of the correlation coefficient for thresholding the network.

**Fig. 1.**
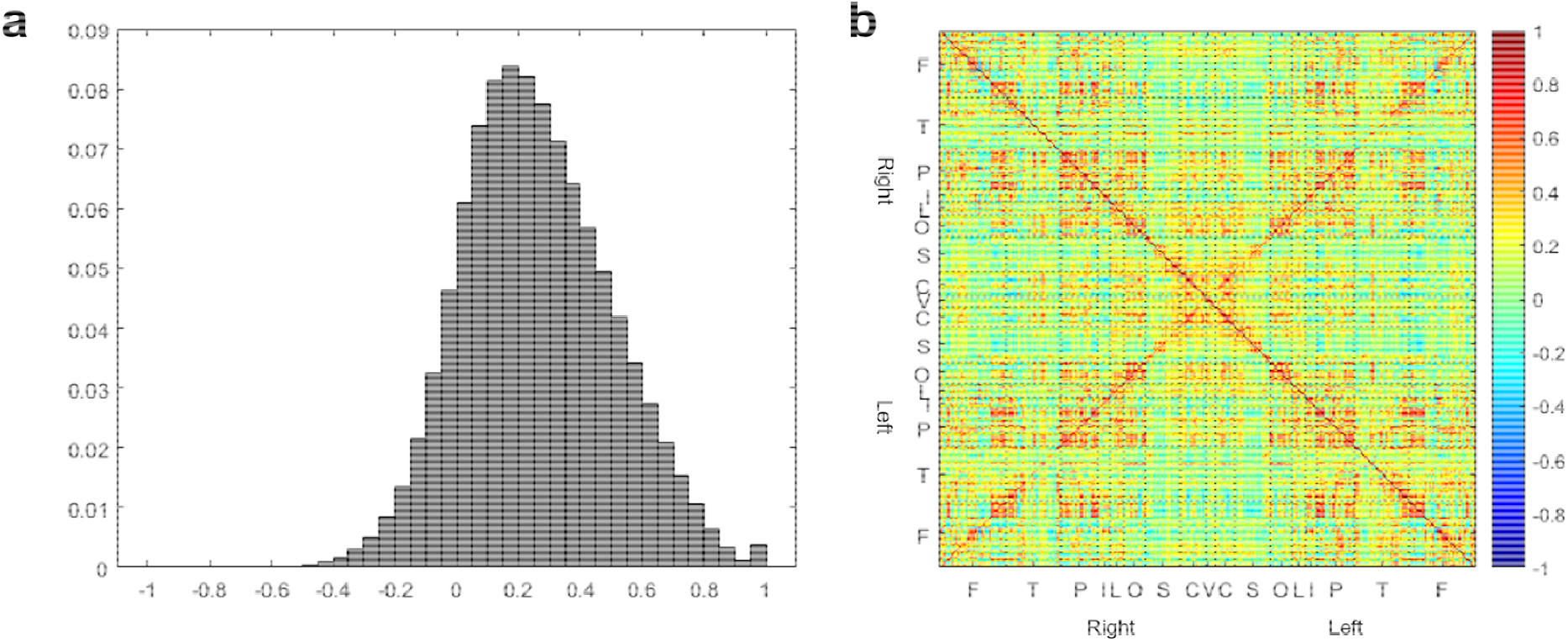
Distributions of time-series correlation of resting-state fMRI signal. **a** Histogram of Pearson’s correlation from all edges human connectome project (HCP) subjects. The Horizontal axis is the value of Pearson’s correlation, while the vertical axis denotes the frequency of the value. The correlation is computed by. A positive correlation between nodes was dominant in healthy adults, while a small portion of edges showed a negative correlation. Overall, 6.70% of edges had a higher absolute value than the threshold and considered as connected. **b** A matrix view of one representative subject. Symbols on each axis denote the anatomic lobe of the nodes. F = Frontal, T = Temporal, P = Parietal, I = insula, L = Limbic, O = Occipital, S = Subcortical, C = Cerebellar, V = Vermis.

### Determining the threshold of correlation for binary network composition

To form the binarized adjacency matrix for each fMRI image, we applied various threshold values to correlation matrix in order to choose binary network that satisfies scale-free property. The chosen threshold yielded degree distribution which fit well to linear plot on log-log relations between degree and degree frequency and included most of the nodes within the largest connected component of the resulting binary network. After we investigated the degree distribution of the graph and the average size of the largest connected component, we chose the threshold for individuals and finally entire population. When we applied a higher threshold, more nodes without any connection were excluded from the component. On the other hand, when we lowered the threshold, the network lost its scale-freeness, which make the later embedding procedure theoretically inappropriate.

Since we wanted to retain as many nodes as possible in every case, we applied the highest threshold possible while the scale-freeness of the network is valid. These procedures are shown in (**Fig. 2**) for three cases of control subjects from Autism Brain Imaging Data Exchange (ABIDE) II dataset. In a threshold value of 0.35 (**Fig. 2a**), in which most of the nodes are preserved, the degree distribution forms a peaked curve, which violates the scale-freeness of the network. While, in a higher threshold value of 0.45 (**Fig. 2c**), 4.8% of nodes, an average of 13 nodes per network is excluded from the largest component, which is considered inappropriate for a whole-brain analysis. Therefore, after searching the entire population of 180 subjects, the threshold value was determined as 0.40 (**Fig. 2b**). For the human connectome project (HCP) dataset, the threshold value was set to 0.36, which resulted from a similar procedure.

**Fig. 2.**
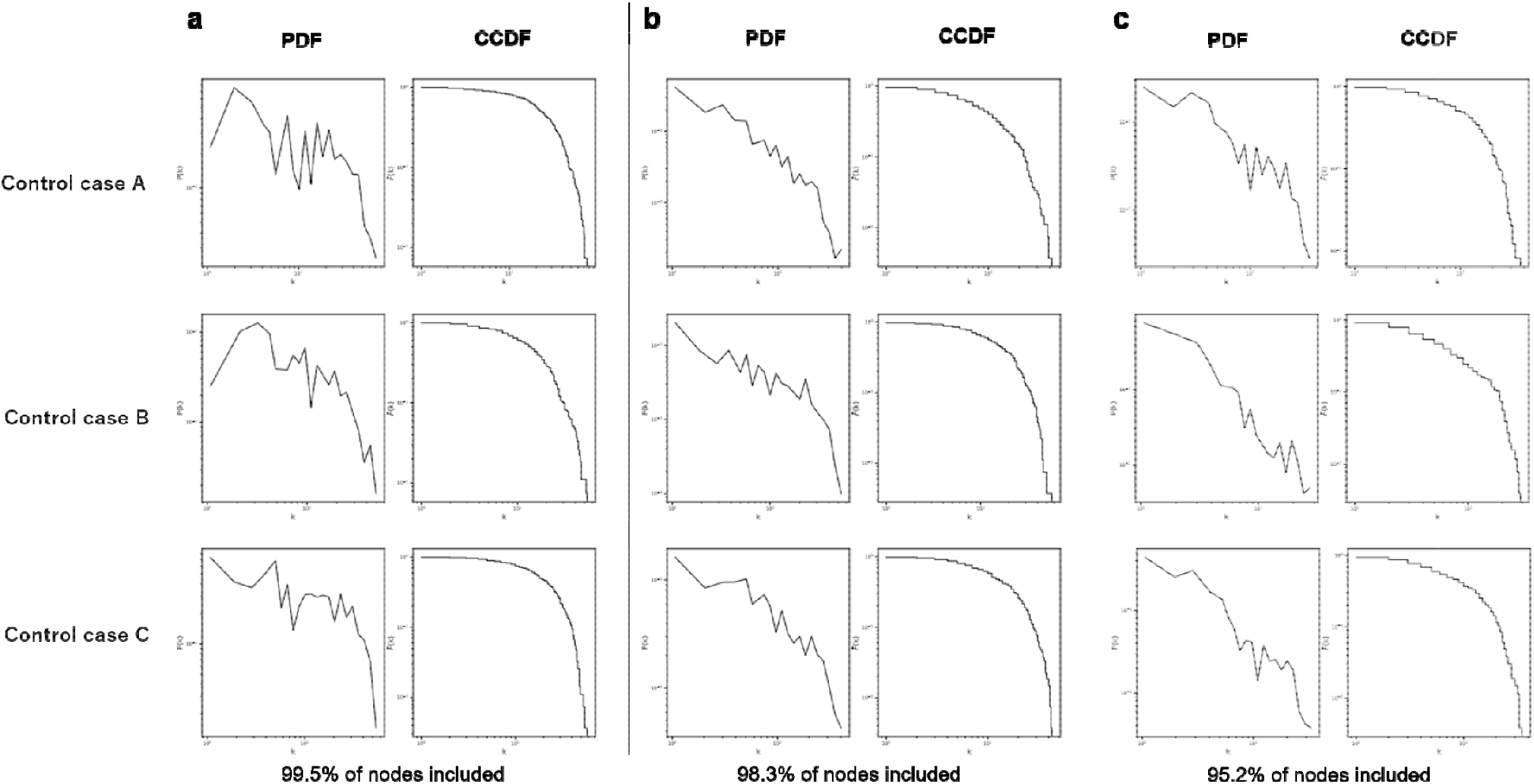
Determining the threshold of correlation for network composition. Probability distribution function (PDF) and complementary cumulative distribution function (CCDF) of degree distribution from three representative subjects from ABIDE II dataset. The threshold is set as **a** 0.35, **b** 0.40, **c** 0.45. **a** In a threshold value of 0.35, in which most of the nodes are preserved, the degree distribution forms a peaked curve, which violates the scale-freeness of the network. **c** While, in a higher threshold value of 0.45, 4.8% of nodes, an average of 13 nodes per network is excluded from the largest component, which is considered inappropriate for a whole-brain analysis. **b** Therefore, the threshold value was determined as 0.40.

### Embedding of the graphs into spaces of various curvatures/dimensions

To find the most natural way of embedding brain graphs into manifolds, we compared the average distortion (D_avg_) and mean average precision (mAP) of embedding between the target manifolds. Ten representative data in the HCP dataset and ABIDE dataset were used for embedding. The target manifolds were hyperbolic, spherical (with unit curvatures, i.e. −1 and 1, respectively), and Euclidean manifolds of 10 and 2 dimensions, plus the single dimension of a spherical manifold 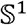 (i.e., circle).

Among the low dimensions, embedding to the 2-dimensional hyperbolic space of 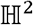 had significantly higher mean average precision (mAP) than embedding to 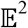 and 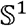. For ABIDE II dataset, embedding to 10-dimensional hyperbolic space of 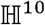 tended to have lower distortion than embedding to 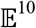. For both datasets, embedding to the 2-dimensional hyperbolic space of 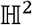 had significantly lower distortion than embedding to 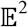 and 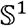 (**Table 1-2, Fig. 3a-b**). Notably, for both datasets, embedding to the 2-dimensional hyperbolic space of 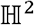 had significantly lower distortion than embedding to 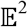 and 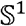 (**Fig. 3c-d**).

**Fig. 3.**
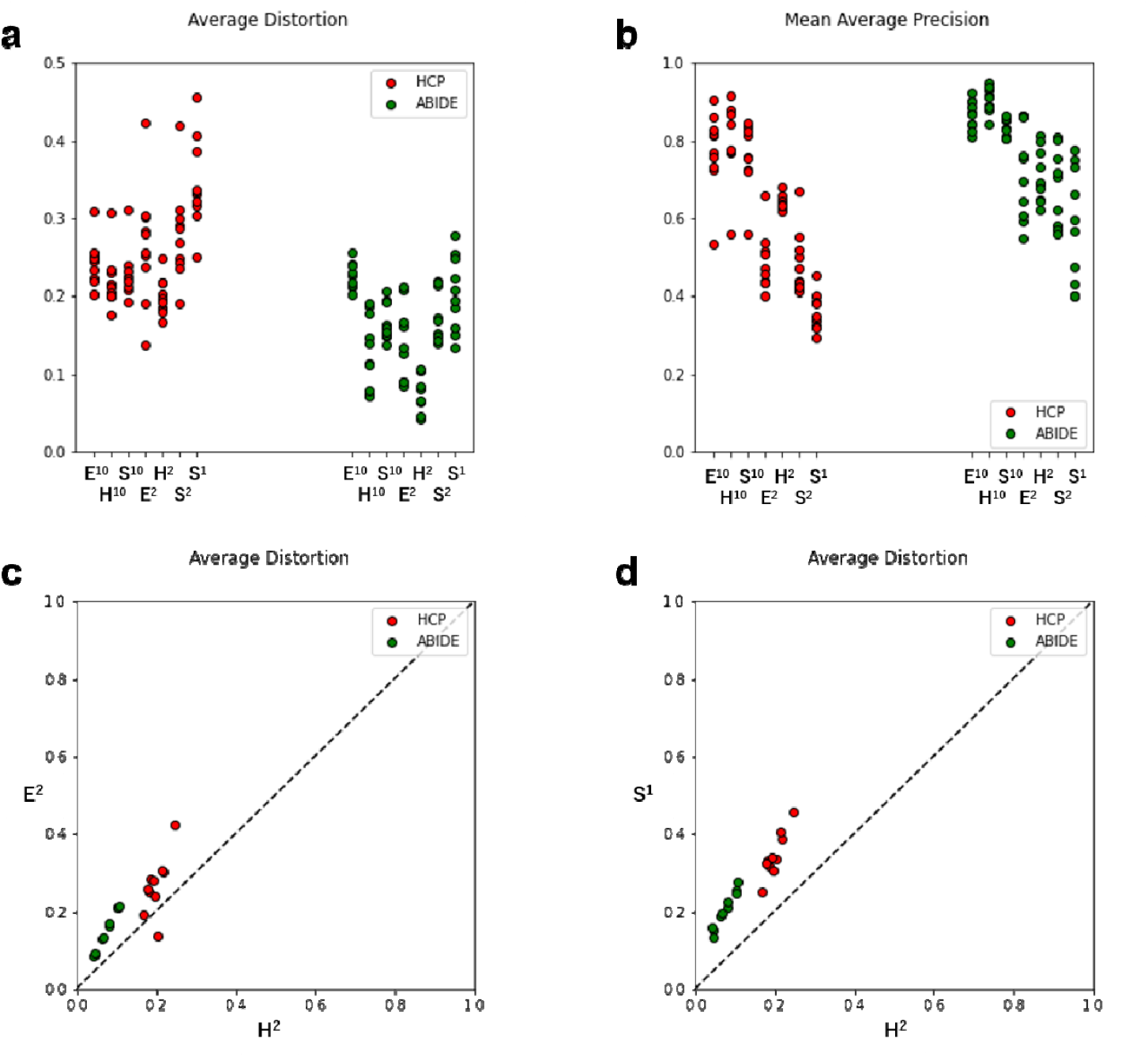
Fidelity measures in embedding onto spaces of various curvatures/dimensions. **a, b** Fidelity measures (average distortion, mean average precision) of embedding for ten representative subjects from HCP and Autism Brain Imaging Data Exchange (ABIDE) II datasets. **c, d** For both of the datasets, embedding to the 2-dimensional hyperbolic space of 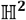 showed lower distortion than embedding to 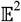 and 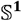.

**Table 1.** Fidelity measures of embedding into various manifolds for HCP dataset. Ten representative cases were used for embedding. For HCP dataset, embedding to the 2-dimensional hyperbolic space of 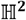 showed higher mean average precision and lower distortion compared to embedding to manifolds 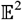 or 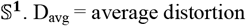. mAP = mean average precision.

| Embedded space | Fidelity measures (Mean ± SD) | |
| --- | --- | --- |
| | $D_{\text{avg}}$ | mAP |
| 10-dimension spaces |  |  |
| $\mathbb{E}^{10}$ | $0.2434 \pm 0.0286$ | $0.7743 \pm 0.1016$ |
| $\mathbb{H}^{10}$ | $0.2181 \pm 0.0351$ | $0.8036 \pm 0.1006$ |
| $\mathbb{S}^{10}$ | $0.2278 \pm 0.0317$ | $0.7675 \pm 0.0874$ |
| 2- and 1-dimension spaces |  |  |
| $\mathbb{E}^2$ | $0.2669 \pm 0.0752^a$ | $0.4854 \pm 0.0740^b$ |
| $\mathbb{H}^2$ | $0.1993 \pm 0.0234^a$ | $0.6404 \pm 0.0175^b$ |
| $\mathbb{S}^2$ | $0.2798 \pm 0.0607^a$ | $0.4834 \pm 0.0803^b$ |
| $\mathbb{S}^1$ | $0.3443 \pm 0.0581^a$ | $0.3575 \pm 0.0482^b$ |

<sup>a</sup> Significantly lower $D_{\text{avg}}$ in $\mathbb{H}^2$ , compared with other results ( $P < 0.05$ ).
<sup>b</sup> Significantly higher mAP in $\mathbb{H}^2$ , compared with other results ( $P < 0.05$ ).

**Table 2.** Fidelity measures of embedding into various manifolds for ABIDE II dataset. For ABIDE II dataset, embedding to 10-dimensional hyperbolic space of 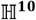 showed lower distortion than embedding to 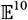. For both of datasets, embedding to the 2-dimensional hyperbolic space of 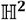 showed lower distortion than embedding to 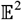 and 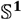.

| Embedded space | Fidelity measures (Mean ± SD) |
| --- | --- |

**Table 2. Fidelity measures of embedding into various manifolds for ABIDE II dataset.**
| | $D_{\text{avg}}$ | mAP |
| --- | --- | --- |
| 10-dimension spaces |  |  |
| $\mathbb{E}^{10}$ | $0.2253 \pm 0.0166^c$ | $0.8674 \pm 0.0372$ |
| $\mathbb{H}^{10}$ | $0.1295 \pm 0.0468^c$ | $0.9030 \pm 0.0318^d$ |
| $\mathbb{S}^{10}$ | $0.1672 \pm 0.0230$ | $0.8316 \pm 0.0237^d$ |
| 2- and 1-dimension spaces |  |  |
| $\mathbb{E}^2$ | $0.1487 \pm 0.0511^e$ | $0.7191 \pm 0.1202^f$ |
| $\mathbb{H}^2$ | $0.0744 \pm 0.0256^e$ | $0.7081 \pm 0.0674^f$ |
| $\mathbb{S}^2$ | $0.1726 \pm 0.0316^e$ | $0.6925 \pm 0.1020$ |
| $\mathbb{S}^1$ | $0.2036 \pm 0.0474^e$ | $0.5793 \pm 0.1477^f$ |

<sup>c</sup> Significantly lower $D_{\text{avg}}$ in $\mathbb{H}^{10}$ , compared with the result in $\mathbb{E}^{10}$ ( $P < 0.05$ ).
<sup>d</sup> Significantly higher mAP in $\mathbb{H}^{10}$ , compared with the result in $\mathbb{S}^{10}$ ( $P < 0.05$ ).
<sup>e</sup> Significantly lower $D_{\text{avg}}$ in $\mathbb{H}^2$ , compared with other results ( $P < 0.05$ ).
<sup>f</sup> Significantly higher mAP in $\mathbb{H}^2$ , compared with the result in $\mathbb{S}^1$ ( $P < 0.05$ ).

### Embedding graphs into the 2-D hyperbolic disc

For each subject, we embedded the largest component of the binary graph to a hyperbolic plane according to the 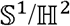 model. Results of the embedding (in the ROI scope) of the representative cases are shown in (**Fig. 4**). Color denotes the anatomic lobe, while the lobes within the right and left cerebral hemispheres and cerebellum are marked with circular, square, and triangular markers, respectively. The results of embedding in the voxel scale are shown in (**Supplementary Fig. 1**).

**Fig. 4.**
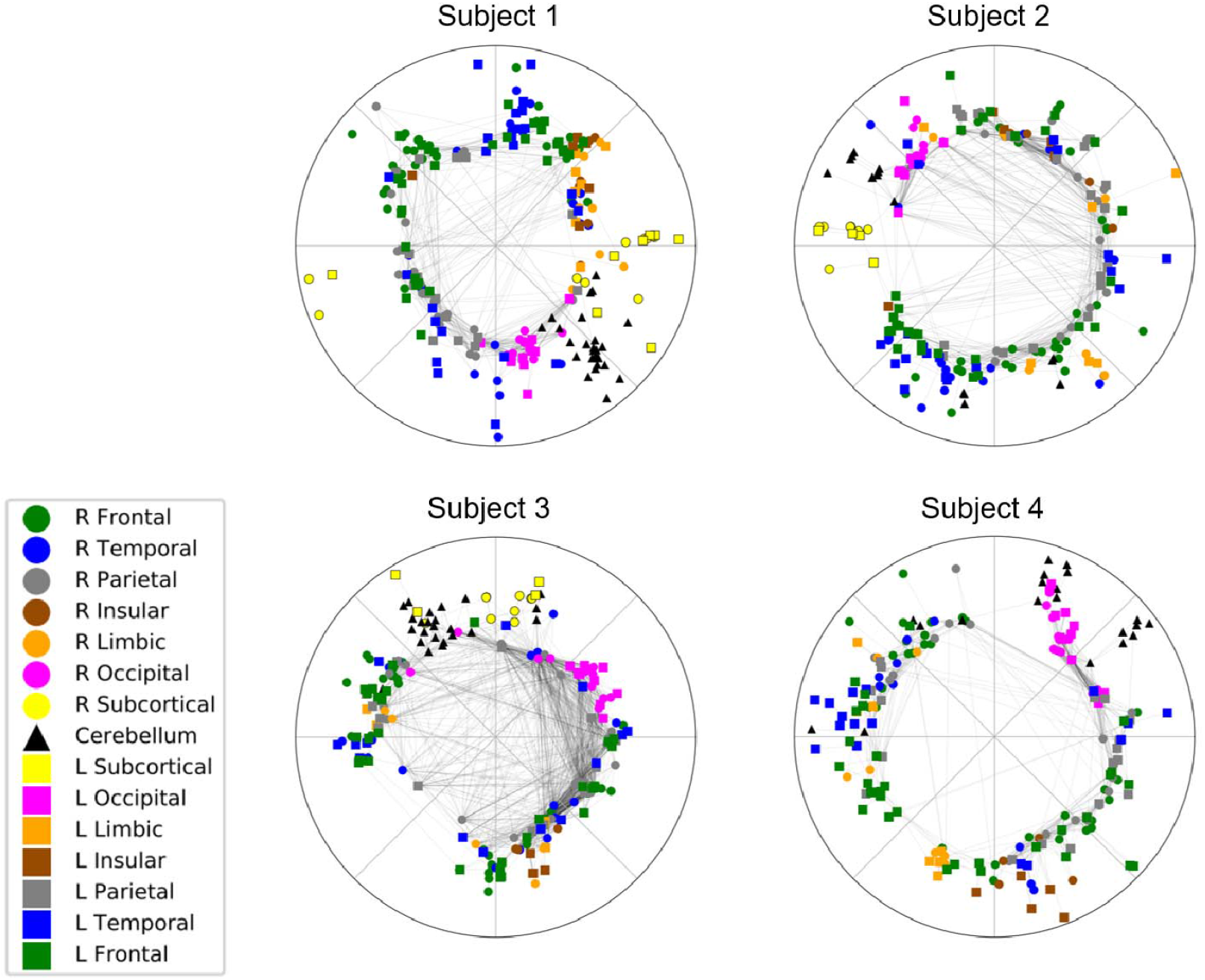
Embedding results by 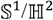 model. The figure demonstrates the result from four representative cases from the HCP dataset. Color denotes the anatomic lobe in which each ROI is located, while the lobes within the right, left cerebral hemisphere, and cerebellum are marked with circle, square and triangular markers, respectively. The embedding result revealed vacant space in the center of the disk. ROIs from the bilateral insula, subcortical, and cerebellum had narrow distributions of angular dimensions. ROIs from larger anatomic lobes (frontal, temporal, parietal) were located with relatively broader distributions. And ROIs from the same lobes of both hemispheres (for example, right and left frontal lobes) had similar distributions of angles.

Every embedding result revealed vacant space in the center of the disk. ROIs from the bilateral insula, subcortical, and cerebellum had narrow angular distributions. ROIs from more large anatomic lobes (frontal, temporal, parietal) were located with relatively broader distributions. And ROIs from the same lobes of contralateral sides (for example, right and left frontal lobes) had similar distributions of angles.

To assess the validity of embedding, we compared the absolute value of original correlation coefficients used for thresholding with connection probability between the edges, which resulted in hyperbolic distances computed on the 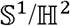 model (**Fig. 5**). The two measures, which are commonly scaled from 0 to 1, tended to show similar patterns for each of the subjects. We discussed the similarity and difference between the two measures in (**Supplementary Note 1**).

**Fig. 5.**
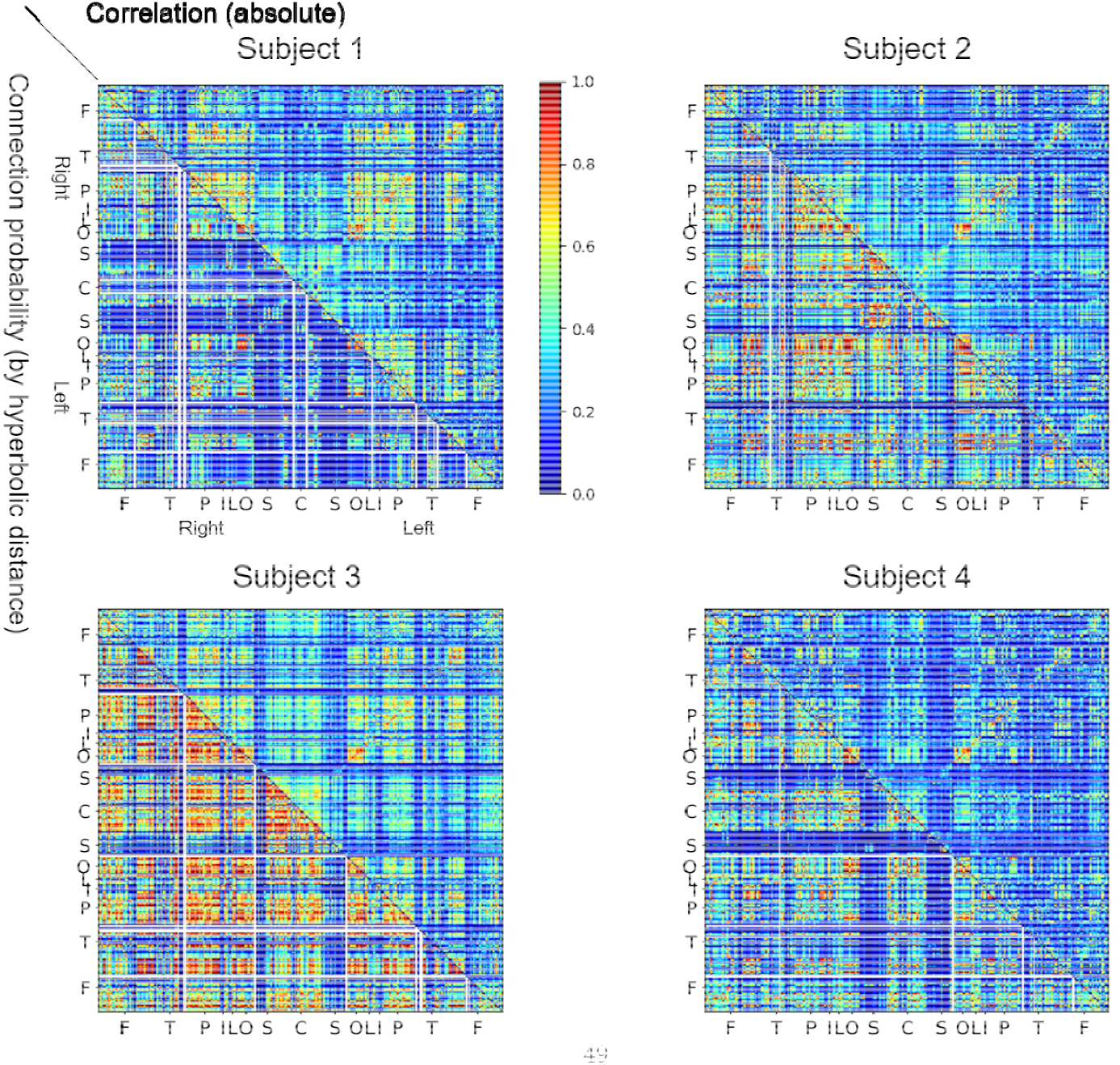
Connection probability versus absolute correlation coefficient. The lower triangle demonstrates the connection probability calculated by 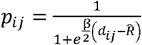 where *d_ij_* is the hyperbolic distance between nodes, β is the clustering coefficient and 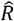 is the outermost radius of the node. The upper triangle denotes the absolute correlation value. The symbols in each axis denote the same anatomic lobe of the nodes as in **Fig. 1**. The two measures tended to show similar patterns for each of the subjects. We discussed the comparison between the two measures in (**Supplementary Note 1**).

To investigate if the embedding results are compatible with other established methods of functional segregation, we made use of network components calculated by independent component analysis (ICA) and plotted the composing voxels of each independent component in colors in the 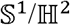 disc (**Supplementary Fig. 2**). To remove the arbitrariness of rotation by the constant angle in the embedded disks, we adjusted the angular position of the embedded disks in order for the mean angle of the visual network (VN) to have a value of 0 (**Supplementary Fig. 3**). However, rotational symmetry should be considered to see the distribution of each component and belonging voxels thereof.

We assessed the reproducibility of the embedding process by measuring the coefficient of variance for the hyperbolic distance matrices in one representative case (**Fig. 6**, **Supplementary Fig. 4**). Most of the nodes had relatively low arbitrariness of position in repeated embedding procedure. Meanwhile, outermost positioned nodes, which represents low popularity, tended to have a position with higher variability (**Supplementary Note 2**).

**Fig. 6.**
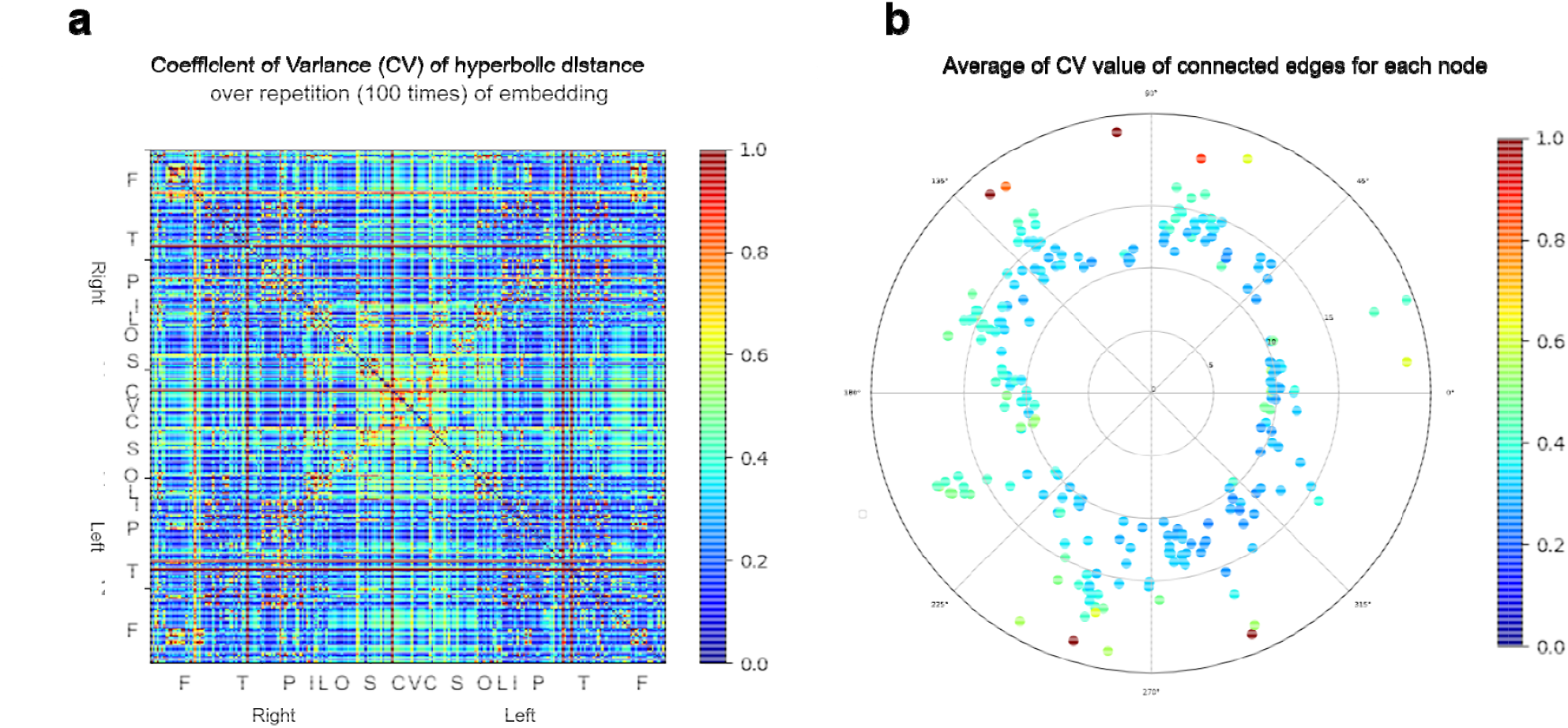
Reproducibility of results (Repetition of the embedding). We repeatedly performed the embedding procedure 100 times, in one representative case. **a** Coefficient of variance (CV) of hyperbolic distances for each edge. The horizontal and vertical axis indicates the indices of vertices embedded in the 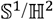 model, which is ordered by lobes. The symbols in each axis denote the same anatomic lobe of the nodes as in **Fig. 1**. **b** Average CV value of connected edges for each node. Blue nodes (low CV) represent nodes with a lower variance of position over embedding. Red nodes (high CV) represent a higher variance. The positions of vertices are positioned based on one sample result of embedding. Most of the nodes have relatively low arbitrariness of position in repeated embedding procedure. In contrast, outermost positioned nodes, which means nodes with low popularity, tended to have a position with higher variability.

### Detection of abnormal functional pathway on hyperbolic disc in the ASD subjects

Among the variable patterns of functional brain networks embedded on 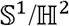 model from ASD group subjects, one subject of autism spectrum disorder showed a longer hyperbolic distance of node pairs connecting the bilateral frontoparietal cortices and bilateral basal ganglia, which represents the cortico-striatal pathway, compared to the control group of 134 healthy young adults (**Fig. 7a**) (*P* < 0.05). This tendency was not prominent when the comparison analysis was conducted on correlation matrices of a subject and 134 controls.

**Fig. 7.**
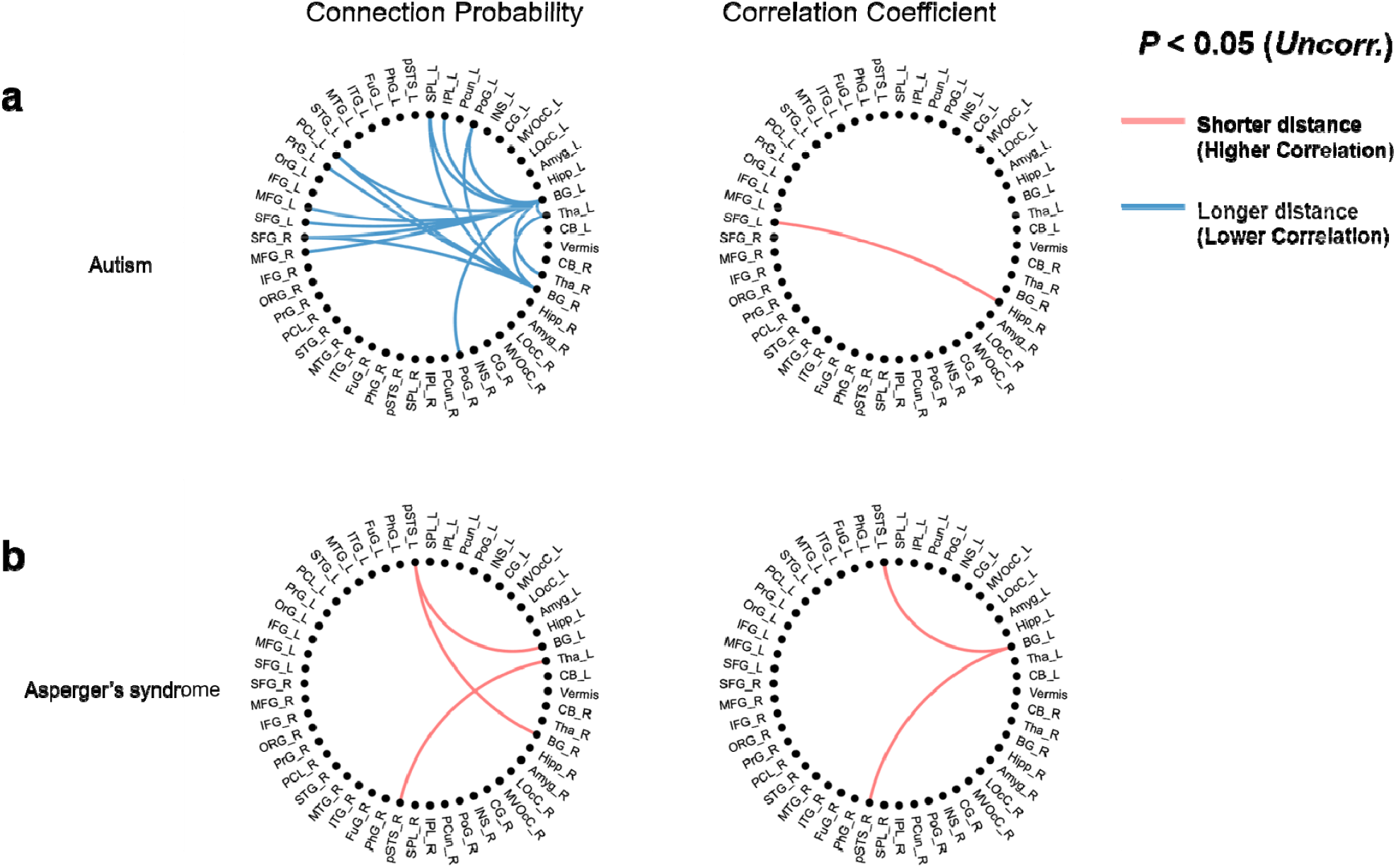
Detection of abnormal pathways in autism subjects. The 274 ROIs were reduced to 51 brain subregions for the sake of clarity of visualization. The lines in the circular diagrams represent the abnormal connection probability or correlation compared to the control group composed of 134 healthy adults. (*Uncorrected P* < 0.05) **a** One subject with autism showed longer hyperbolic distances of edges connecting frontoparietal cortices and bilateral basal ganglia, which represents the cortico-striatal pathway, compared to the normal group. The pattern was not clear in the means of the correlation coefficients. **b** Another subject with Asperger’s syndrome showed shorter hyperbolic distances of edges which connect bilateral posterior superior temporal sulcus (pSTS) and basal ganglia. The correlation study showed a similar pattern. The list of abbreviations in the circular diagrams is noted in (**Supplementary Note 4**).

Notably, a subject with Asperger’s syndrome showed a shorter hyperbolic distance of the edges of node pairs, which connect bilateral posterior superior temporal sulcus (pSTS) and basal ganglia, and the correlation study showed a similar pattern (**Fig. 7b**) (*P* < 0.05).

## Discussion

Neuroscientific work in the past decades has provided the perspective that the brain is a complex network that is composed of interactions between regions^17^, and rs-fMRI has emerged as an efficient tool in exploring the connection^18^. However, it still remains unclear how each region of the brain is connected functionally in a multiplex way and organized in higher order to perform its complicated function at rest with activation whose amplitude was smaller than when related with tasks. Preceding work by Allard et al.^7,19^ implemented hyperbolic geometry model to enlighten this question by providing a map of structural connectomes, while exploiting this geometry’s expansive space able to contain brain connectome’s high-order and high-dimensional correlational structure. One of the major advantages of this model is that it implements 2-dimensional hyperbolic space to embed the network, which is makes it easy to visualize brain connectomes and to understand of how the brain regions are working in a multiplex connected fashion.

Meanwhile, not much work has attempted to yield deep insight of this subtle correlative works of multiple nodes of an individual brain from the norms made of many subjects as represented by an individual functional map to define an unheralded disease-specific anomaly of a single affected subject. Since many neurologic disorders are classified as a group of similar clinical manifestations^20^ with unclear neurologic pathophysiology, changing the scope into connectomes of affected individuals might significantly help in further classifying the disease entities and provide information in the treatment of disease. Furthermore, this work inherits the advantage of the preceding work by Allard et al., that the resulting object is represented in 2-dimensional space both for visualization and for further analysis to disclose which internodal connections are deviated from the norms in these hyperbolic discs.

### Composition of the network

To compose a framework of the functional brain network, we implemented inter-region time-series correlations of rs-fMRI, which grants a stable measure of the resting brain^21^. We applied the *absolute value* of Pearson’s correlation between brain regions, which regards the two brain regions with anticorrelated time series as connected, as same as the correlation of the same strength. Anti-correlation between specific brain regions has a role in organizing functional brain architectures^22–24^. Then, we thresholded the matrix to obtain a binary graph for each network. In this study, we decided to apply the same threshold among patients in a dataset for comparability.

Since only the one largest component can be used to be embedded in the model, we needed to retain as many nodes as possible. If we apply the higher threshold value, we drop out more nodes from the network, which makes it difficult to compare between individuals and establish the difference clinically meaningful from the embedding of the whole-brain networks. On the other hand, if we apply a too low threshold value, the binary networks become denser with almost every edge connected, and the assumption of scale-freeness is violated as we perform maximum likelihood estimation for embedding the functional brain connectomes to the hyperbolic disc using the 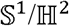 model embedding. Thus, we composed the sparsest network while maintaining most of the brain regions in its largest component. We assume that we could remove noise and/or artifacts in the observational data and also emphasized the multi-dimensional tree-like internodal correlational structure of the functional brain networks.

### Scale-freeness of the network

The degree distributions of binarized brain networks showed patterns of straight downward lines in the log-log scale graph rather than concave or convex lines, which reveals that these networks have power-law degree sequences, which is a feature of a scale-free network^2^.

That a network is a scale-free network means that the structure of the network has a similar structure independent of the scale of observation^25^. In detail, most nodes have a small number of connections, while some nodes have a large number of functional connections with other nodes. This indicates some of the topological features of the functional brain network, such as the hierarchical organization of nodes^26^, as its structural counterpart^27^. Heterogeneity of degree distribution implies various models including hyperbolic one and we adopted hyperbolic model in this investigation.

### The geometry of the brain network

Real-world networks can be embedded in a physical space corresponding to their structure and organization^28,29^. In this study, the architecture of the high-order high-dimensional internodal correlational structure of functional brain network, of which structural counterpart lies in 3-D Euclidean space, was investigated by comparing and computing the fidelity measures of embedding in manifolds of positive (spherical), negative (hyperbolic) and zero (Euclidean) curvatures. Our result showed the better quality of embedding in 2-D hyperbolic spaces compared with the 2-D Euclidean or 1-D spherical spaces, which justifies further embedding of the network into the 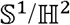 model, which is of 2-D hyperbolic space. Low distortion, global measure of fidelity from the original correlational structure, meant hyperbolic disc representation can be used as surrogate of the original functional brain network. Higher precision, local measure of fidelity, meant again the same.

Space of hyperbolic geometry is in fact, just the place we live in, as indicated by the special relativity with multi-layered, hierarchical and multiplex-correlated way. The most characteristic feature about the hyperbolic space against other types of space is that it best represents the data with hierarchical structure^30^, as in deep neural network implemented in deep learning^31^. Prior research about embedding into non-Euclidean spaces also reports the hyperbolic space as the matching geometry to embed tree-like structures, employing the least distortion^32,33^. This reveals the tree-like nature of the network, follows from the nature of the hyperbolic space that the geodesic passes path near the origin, as the shortest path in the tree graph passes their common parent node. It should be reminded that functional brain networks form tree structure, not in a 3-dimensional space of real life, but in a higher-dimensional space derived from the correlations of nodes during 5 minutes (ABIDE) or 15 minutes (HCP) of rs-fMRI acquisition based on the stationarity assumption. The hyperbolic geometry explains that the graphs, derived from this internodal functional brain connectomes, that represent the brain networks exhibit hierarchical and tree-like organization, which is suggested in prior research of humans or other animals^34,35^.

### Hyperbolic disc representation of the network

Since it has been known that many real-world networks have hyperbolic properties, many investigators have proposed algorithms for figuring out how to generate geometric objects that are most likely to generate the given network^36–39^. The present study makes use of *Mercator*^15^, using Laplace eigenmaps (LE) for the reduction of dimension and maximum likelihood estimation (MLE) techniques for acquiring the most appropriate geometric object representing the original network^38^. By this means, we could visualize the functional brain network in the hyperbolic disc with the conformal structural properties with the real-world network^14,40^. Note that the two embeddings we used for testing fidelity of embedding on the spaces with various curvatures/dimensions and for final hyperbolic disc embedding represent two different procedures. In the former task for testing embedding spaces, we used the learning procedure by minimizing the loss function^32^, while in the latter task of 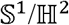 model embedding, we used *Mercator*^15^ for reduction of dimension and maximizing likelihood for the geometric object to generate the original binary graph^14^.

Embedding procedure of the binary binary graph onto the hyperbolic disc according to 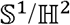 model provides the hyperbolic distance as a measure of how the two brain nodes are functionally similar, i.e., similar correlational structures derived from time-series of node pairs. As an effective distance^7^, this combines factors that determine the connection probability between nodes (which are related as in Eq. (6) in the methods). As a result, this procedure effectively visualizes the network to give a quantitative comparison of the functional brain network among individuals.

The geometric meaning of the 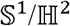 model is that the popularity and similarity of the nodes are represented by the radial and angular coordinate of the 2-D hyperbolic space, respectively^14^. The basic assumption is that the closer nodes are more likely to be connected. In other words, the short hyperbolic distance between the two nodes correlates with a high probability of connection in the generated network, and this process of embedding helps us to understand the topology of the network and growth dynamics for the system^41^.

The radial representation of the embedded nodes of the network represents is hub characteristics or popularity. That is, nodes with higher degrees are located closer to the center of the disk. From this perspective, we point out that in all of our embedding results, centers of embedded disks were vacant. This implies that the functional brain network does not have any nodes with dominant high popularity implying that functional brain networks have a decentralized information flow.

Previous similar studies performed with other kinds of real networks showed variable patterns. The network of world trade atlas in 2013^13^ showed two global hubs remarkably close to the center of the disk, named “USA” and “China.” These two nodes work as the global hub, which generally has a high connection probability with any other nodes from the whole network. On the other hand, the embedded map of internet connection^12^ did not show heavily centered nodes, but several locally centered nodes were seen, which might function as a local hub of the system. Finally, the map of the structural brain network^7^ showed a relatively uniform distribution of radial coordinate without noticeably centered node.

The mapping result of the functional brain network is most similar to the result from the structural network, which indicates that there is no definite global hub connected with the vast substructures of the network. This might result from the fact that both structural and functional brain network is based on the real neuronal network of the brain, of which the number of connection (i. e. degree of the node) is limited by physical constraints. Note that in the former two examples, the number of connections for each node is not physically limited. This does not mean that the hierarchy does not exist: the nodes have variation in their radial coordinates, which means it does exist in the functional brain network.

From the tendency that nodes from the same anatomic lobe tend to cluster in a similar sections on the angular coordinates, we could address a certain degree of neuroanatomical relevance, which is reported from previous literature on functional brain networks^42^. The regions involved in sensory or motor function such as subcortical regions, occipital lobes and cerebellum showed relatively strong concentration in narrow sections of angular coordinates, while the association cortices such as frontal, temporal and parietal regions were broadly distributed in angle.

Another notable feature is that nodes from the same lobes of both hemispheres, which are also denoted as the homotopic area, tend to cluster in similar angular coordinates, wherever they are located. This is markedly different from the map of the structural brain network, in which the different hemispheres showed separation on angular coordinates. As homotopic connection in the brain is reported by prior research^43–45^, our results suggest a stronger homotopic connection in functional connectivity, more than indicated by structural connection.

Note that the set of positions of nodes has symmetry upon reflection and rotation^46^ and branch permutation invariance^47^, which inevitably arise when we adopt hyperbolic embedding of any kind, Poincare or other hyperbolic ones. These are key features to consider when one compares a (group of) disc to another, and implementing the hyperbolic distance between two nodes, which is a *relative* measure, resolves the reflection and rotational symmetry issue. Since hyperbolic distance is an essential measure of this 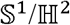 model because it is the single argument that determines connection probability between two nodes of the disc, we could also assure the reproducibility of the result in terms of hyperbolic distance.

### Abnormality of functional pathways in the diseased subjects

The autism spectrum disorder (ASD) is a diagnostic group of neurodevelopmental disorders with deficits in communication and social interaction and repetitive, restrictive behaviors^48^. A vast number of studies in neuroscience have accumulated sufficient data to make clear that ASD is a disorder of abnormal connectivity of neural pathways^49–52^, in spite of its complex and heterogeneous nature^53^.

We assessed the clinical applicability of our method by detecting abnormalities of the network in the ASD subject compared with the control group. While the abnormality of network on hyperbolic discs among subjects was variable, some subjects showed results congruent with clinical context and literature reports. The striatum is pointed to as the key of the pathophysiology of autism^54,55^, and one of the subjects showed abnormal hyperbolic distance in pathways connecting the frontoparietal cerebral cortex and striatum (**Fig. 7a**).

Asperger’s syndrome, a high-functioning form of autism, is characterized by a higher intelligence and better-than-average verbal skills, but impaired nonverbal communication and social interaction, with restrictive and repetitive behaviors as in classic autistic patients^56,57^. Though our study has presented the case of a single subject with Asperger’s syndrome, the results showed an abnormal connection pattern of edges connecting the bilateral posterior superior temporal sulcus (pSTS) and subcortical regions compatible with the literature reports^58,59^. The bilateral pSTS is known to be associated with social interactions, and some prior literature reports its relevance with ASD. Our result suggests the abnormal connection of pSTS, which is consistent with impairment of social interaction in high-functioning autism patients, while the cortico-striatal pathway was relatively intact.

There are several limitations of this analysis. First, in the composition of the network, we used the absolute value of the correlation coefficient, which regards the anticorrelation between brain regions same as the same degree of correlation. Since the anticorrelation between brain regions is believed to have a distinct nature^60,61^, we should consider multigraphs to take both types of correlation into account in the future study. Second, the embedding we performed in searching for the best fit manifold among spaces with various curvatures/dimensions is not the same as we have done finally to represent functional brain networks on hyperbolic discs according to 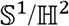 model, and followingly, findings from the former investigation of comparison between various space embeddings might not guarantee the highest quality of finally used hyperbolic disc embedding over any other choices. Third, we did not perform the multiple comparison adjustment in detecting the abnormality of the network, which reduces the statistical significance.

In conclusion, in this work, we aimed to find the most appropriate geometry for functional brain networks based on inter-regional time series correlation of rs-fMRI, and we embedded the networks onto 2-D hyperbolic discs of the 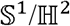 model. The results demonstrated the absence of global hub, homotopic functional coherence, and a certain degree of functional-anatomical relevance and yielded a reliable parameter of hyperbolic distances on this hyperbolic disc. When this method was applied to the diseased subjects, we detected abnormal pathways by comparing internodal hyperbolic distances. This leads us to a new method of reproducible visualization of functional brain networks with high fidelity from correlational functional brain networks and anomaly detection of internodal pathways in the brain networks of diseased subjects.

## Acknowledgments

This research was supported by the National Research Foundation of Korea (NRF) grant funded by the Korean Government (MSIT) (No. 2017R1A5A1015626, No. 2017M3C7A1048079 and No. 2020R1A2C2101069) and NRF grant funded by the Korean Government (No. 2017R1D1A1B03032037). We deeply appreciate Dr. M. Ángeles Serrano (Universitat de Barcelona) for helpful advice on the methodology, and Dr. Seonhee Lim (Seoul National University) for her help and essential advice in constructing the theoretical basis of the research and further comments that greatly improved the manuscript.

## Author contributions

W. W., H. K., and D. S. L. designed this study. H. K. designed the methods of diseased subejcts. W. W. and H. K. performed data collection and analysis. W. W., S. H. and D. S. L. conducted the clinical interpretation. All authors interpreted data, drafted, and edited the manuscript.

## Methods

### Subjects

#### Human Connectome Project (HCP) dataset

To investigate the general characteristics of the functional brain network, we used the rs-fMRI datasets from healthy adults of the Human Connectome Project (HCP) S1200 release. Participants were between 22 and 35 years of age at the time of recruitment and did not have any documented history of psychiatric, neurological, or medical disorders known to influence brain function. A more detailed description of the inclusion and exclusion criteria for HCP is shown in the literature^62^. The sample of participants included one hundred and eighty subjects (Mean age = 29.1, SD = 3.4), with 76 male and 104 female subjects.

#### Autism Brain Imaging Data Exchange II (ABIDE II) dataset

For the investigation of the characteristic of autistic spectrum disorder (ASD) diseased subject, we made use of 247 resting-state fMRI datasets from Autism Brain Imaging Data Exchange II (ABIDE II) datasets, 113 from diseased subjects, and 134 from the control group. We selected data from the same age group as that of HCP datasets, whose age ranged between 20 and 35 years of age in the control group (Mean age = 26.8, SD = 3.8), and 20 to 39 years of age in the diseased group (Mean age = 27.9, SD = 4.9). A more detailed description of the subject information and image acquisition protocol is described in the literature^63^.

### Functional connectivity analysis

We parcellated the whole brain in two scopes; a) 274 regions of interest (ROIs) using the human Brainnetome Atlas^64^. All of the 274 sub-ROIs were included in the analysis. Detailed information on ROIs, including full name and abbreviations, are available in (https://atlas.brainnetome.org/). 2) cubic isotropic voxels of 6 × 6 × 6 mm^3^ size, the total number of 5,937 voxels. The voxel-scope analysis was restricted to ten randomly selected cases of the HCP dataset due to long computation time.

Spontaneous fluctuations were characterized by the variance of the population σ^2^(*X*) of the BOLD signal of fMRI time-series X^59^. For the BOLD-fMRI time-series X = (X_1_,…, X_N_) of a given ROI, the variance was computed by the sample variance 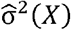

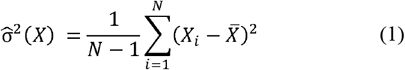

in Eq. (1), 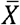 denotes the sample mean of *X*. Functional connectivity was assessed by the Pearson correlation coefficient ρ:

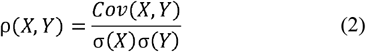

where (*X, Y*) stands for the BOLD-fMRI time-series. For a pair of BOLD-fMRI time-series X = (X_1_,…, X_N_) and Y = (Y_1_,…, Y_N_), 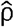 was estimated by the sample Pearson correlation coefficient 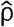:

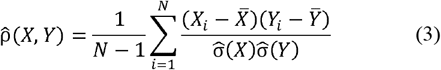

From the Pearson correlation coefficients, we obtained a square matrix of Pearson correlation coefficient X for each of the subjects. The connectivity matrix that consists of absolute values of both positive and negative correlation coefficients was used to establish a binary graph from the network. We tried multiple threshold values for each group analysis and compared the degree distribution and the number of nodes belonging to the largest connected component. We determined fixed threshold values (0.36 for the HCP dataset and 0.40 for ABIDE II dataset) by considering both the scale-freeness and the maximal inclusion of nodes in the chosen brain network. Consequently, we constructed an unweighted, undirected graph for each subject by applying the threshold to the coefficient matrix.

### Embedding the network into manifolds of various curvatures and dimensions

An embedding of the network is assigning nodes to representative low-dimensional space, which effectively preserves the network structure^65^. The analytical meaning of embedding is the mapping *f*:*U*→*V* for spaces *U, V* with distances *d_U_, d_V_*. We can measure the quality of embeddings with fidelity measures. The standard metric for graph embeddings is distortion *D* ^33^. For an *n*-point embedding,

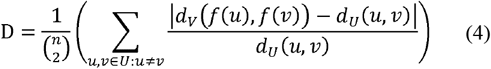

The best distortion is *D*(*f*) = 0, preserving the edge lengths exactly. Also, note that *D*(*f*) can be larger than 1. This is a *global* metric, as it depends directly on the value of hyperbolic distances rather than the local relationships, or ranks, between distances.

Preceding work by Nickel & Kiela et al.^66^ suggests mean average precision (mAP) for another measure. For a graph *G* = (*V, E*), let *a* ∈ *V* have neighborhood 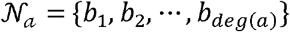, where *deg*(*a*) denotes the degree of *a*. In the embedding *f*, consider the points closest to *f*(*a*), and define *R_a,b_i__* be the smallest set of such points that contains *b_i_*, i.e., *R_a,b_i__* is the smallest set of nearest points required to retrieve the *i*th neighbor of *a* in *f* ^67^.

Then, the mAP is defined to be

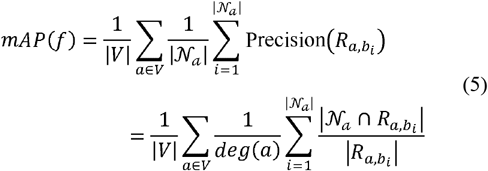

Note that mAP(*f*) ≤ 1. And the equal sign is the best case that preserves the rank of distances between the nodes. The mAP is not concerned with the exact value of underlying distances but only the relative ranks between the distances of immediate neighbors. This is a *local* metric.

To compute the quality of embeddings, we implemented the method proposed by Chami et al.^28^ to optimize the placement of points using an auxiliary loss function^a^.

### *Embedding onto hyperbolic discs of* 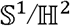 *model*

The binary graph from each patient was then embedded into latent hyperbolic geometry of 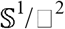 geometric network model. In the model, the connection probability between two nodes *i* and *j* is determined by the hyperbolic distance between two nodes:

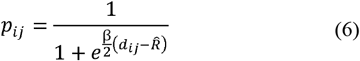
where *d_ij_* is the hyperbolic distance, which has a good approximation:

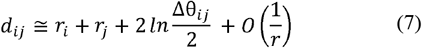
β is the clustering coefficient of network and 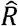 is the outermost radial coordinate among the embedded nodes. We described the detail of 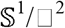 geometric model in the (**Supplementary Note 3**). To find out the most appropriate geometric object (i.e., a hyperbolic disc) that is most likely to generate the binary graph we made, we implemented a software named *Mercator*^b^, introduced by García-Pérez et al.^15^, which assumed that the structure of networks could be described by the 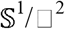 geometric model. We embedded an undirected, unweighted adjacency matrix for each subject onto a hyperbolic disc. In other words, we searched a set of angular coordinates for *N* nodes (r_1_, θ_1_), (r_2_, θ_2_),⋯(r_*N*_, θ_*N*_) and the clustering coefficient β on a hyperbolic disc.

### Comparison with ICA-driven results

To investigate the compatibility of the embedding with other established methods, resting-state networks determined from group independent component analysis (ICA)^68^ were plotted on the individual hyperbolic discs for the voxel data. Each independent component (IC) was thresholded by a z-score larger than 6. As ROI-derived correlation matrix was embedded onto 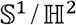 modeled hyperbolic discs, voxel-derived correlation matrix on rs-fMRI was embedded onto hyperbolic discs and colored for the lobes used in ROI-lobe coloring (Supplementary Fig. 1) and also each independent component such as default mode network (DMN), salience network (SN), visual network (VN) and others in voxel-independent component (IC) coloring (**Supplementary Fig. 2**).

### Reproducibility of embedding in an individual

We computed the hyperbolic distance between ROIs over multiple performances of embedding for one representative case and computed the coefficient of variance (CV) for distance analog, which is correlated with connection probability,

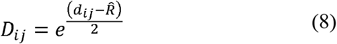

which appears denominator of the right term at Eq. (6) when we assume β equals 1 and is a simple increasing function *d_ij_*, for each pair of nodes. We averaged connection probability of edges of a certain node with every other nodes and the averages were assumed to be the average connection probability of this node and on the repeated embedding 100 times, the CV was calculated for this node. Each node has CV and this CV was plotted on the hyperbolic disc of any one embedded disc among 100 embedded discs of repetition. An example was shown in (**Fig. 6b**).

### Detection of abnormality of hyperbolic distance in the diseased subjects compared with controls

To find the abnormal pathway of the functional brain network in ASD group subjects, we used the ABIDE II dataset and compared the distance matrices resulting from the 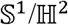 model embedding, according to the following process.

After embedding the individual networks according to the 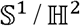 hyperbolic embedding in the ROI scale, we computed the hyperbolic distance *d_ij_* determined by Eq. (7) between each pair of ROIs for each individual subject in disease group and the control subjects, i.e., individual to group comparison. We set the distribution of hyperbolic distances for each edge in the control group dataset. For each ASD group subject, we considered significant edges with higher/lower hyperbolic distances seen on the 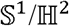 models.

To compare the results with those from the traditional correlation study, we conducted a similar comparison by lower/higher correlation values between an individual and the controls group. We selected edges of which hyperbolic distance was located in upper or lower 2.5% of the distribution. In the control group, hyperbolic distance and correlation value for each edge were tested for normality (P<0.05). Multiple comparison correction was not performed.

## Footnotes

a Available at https://github.com/HazyResearch/hyperbolics

b Available at https://github.com/networkgeometry/mercator

## References

1 Betzel, R. F. et al. Changes in structural and functional connectivity among resting-state networks across the human lifespan. Neuroimage 102, 345–357 (2014).

2 Voitalov, I., van der Hoorn, P., van der Hofstad, R. & Krioukov, D. Scale-free networks well done. Physical Review Research 1, 033034 (2019).

3 Meunier, D., Lambiotte, R. & Bullmore, E. T. Modular and hierarchically modular organization of brain networks. NeuroImage 4, 200 (2010).

4 Bassett, D. S. & Bullmore, E. T. Small-world brain networks revisited. The Neuroscientist 23, 499–516 (2017).

5 Gastner, M. T. & Ódor, G. The topology of large Open Connectome networks for the human brain. Scientific Reports 6, 1–11 (2016).

6 Tadic, B., Andjelković, M. & Melnik, R. Functional geometry of human connectomes. Scientific Reports 9, 1–12 (2019).

7 Allard, A. & Serrano, M. Á. Navigable maps of structural brain networks across species. PLoS Computational Biology 16, e1007584 (2020).

8 Stam, C. et al. The relation between structural and functional connectivity patterns in complex brain networks. International Journal of Psychophysiology 103, 149–160 (2016).

9 Markram, H. et al. Reconstruction and simulation of neocortical microcircuitry. Cell 163, 456–492 (2015).

10 Sporns, O. Structure and function of complex brain networks. Dialogues in Clinical Neuroscience 15, 247 (2013).

11 Clauset, A., Moore, C. & Newman, M. E. Hierarchical structure and the prediction of missing links in networks. Nature 453, 98–101 (2008).

12 Boguná, M., Papadopoulos, F. & Krioukov, D. Sustaining the internet with hyperbolic mapping. Nature Communications 1, 1–8 (2010).

13 García-Pérez, G., Boguñá, M., Allard, A. & Serrano, M. Á. The hidden hyperbolic geometry of international trade: World Trade Atlas 1870-2013. Scientific Reports 6, 1–10 (2016).

14 Papadopoulos, F., Kitsak, M., Serrano, M. Á., Boguná, M. & Krioukov, D. Popularity versus similarity in growing networks. Nature 489, 537–540 (2012).

15 García-Pérez, G., Allard, A., Serrano, M. Á. & Boguñá, M. Mercator: uncovering faithful hyperbolic embeddings of complex networks. New Journal of Physics 21, 123033 (2019).

16 Serrano, M. A., Krioukov, D. & Boguná, M. Self-similarity of complex networks and hidden metric spaces. Physical review letters 100, 078701 (2008).

17 Sporns, O. The human connectome: a complex network. Annals of the New York Academy of Sciences 1224, 109–125 (2011).

18 Van Den Heuvel, M. P. & Pol, H. E. H. Exploring the brain network: a review on resting-state fMRI functional connectivity. European Neuropsychopharmacology 20, 519–534 (2010).

19 Allard, A., Serrano, M. Á., García-Pérez, G. & Boguñá, M. The geometric nature of weights in real complex networks. Nature communications 8, 1–8 (2017).

20 Thapar, A., Cooper, M. & Rutter, M. Neurodevelopmental disorders. The Lancet Psychiatry 4, 339–346 (2017).

21 Greicius, M. D., Krasnow, B., Reiss, A. L. & Menon, V. Functional connectivity in the resting brain: a network analysis of the default mode hypothesis. Proceedings of the National Academy of Sciences 100, 253–258 (2003).

22 Kucyi, A. et al. Intracranial electrophysiology reveals reproducible intrinsic functional connectivity within human brain networks. Journal of Neuroscience 38, 4230–4242 (2018).

23 Keller, J. B. et al. Resting-state anticorrelations between medial and lateral prefrontal cortex: association with working memory, aging, and individual differences. Cortex 64, 271–280 (2015).

24 De Havas, J. A., Parimal, S., Soon, C. S. & Chee, M. W. Sleep deprivation reduces default mode network connectivity and anti-correlation during rest and task performance. Neuroimage 59, 1745–1751 (2012).

25 Barabási, A.-L. & Bonabeau, E. Scale-free networks. Scientific American 288, 60–69 (2003).

26 Sporns, O., Chialvo, D. R., Kaiser, M. & Hilgetag, C. C. Organization, development and function of complex brain networks. Trends in Cognitive Sciences 8, 418–425 (2004).

27 Faskowitz, J., Yan, X., Zuo, X.-N. & Sporns, O. Weighted stochastic block models of the human connectome across the life span. Scientific Reports 8, 1–16 (2018).

28 Chami, I., Wolf, A., Sala, F. & Ré, C. Low-dimensional knowledge graph embeddings via hyperbolic rotations. in Graph Representation Learning NeurIPS 2019 Workshop.

29 Smith, A. L., Asta, D. M. & Calder, C. A. The geometry of continuous latent space models for network data. Statistical Science: a Review Journal of the Institute of Mathematical Statistics 34, 428 (2019).

30 Lazcano, D., Fredes, N. & Creixell, W. Hyperbolic Generative Adversarial Network. arXiv preprint arXiv:.05567 (2021).

31 Peng, W., Varanka, T., Mostafa, A., Shi, H. & Zhao, G. Hyperbolic Deep Neural Networks: A Survey. arXiv preprint arXiv:.04562 (2021).

32 Gu, A., Sala, F., Gunel, B. & Ré, C. Learning mixed-curvature representations in product spaces. in International Conference on Learning Representations.

33 Sala, F., De Sa, C., Gu, A. & Ré, C. Representation tradeoffs for hyperbolic embeddings. in International Conference on Machine Learning. 4460–4469 (PMLR).

34 Telesford, Q. K., Simpson, S. L., Burdette, J. H., Hayasaka, S. & Laurienti, P. J. The brain as a complex system: using network science as a tool for understanding the brain. Brain Connectivity 1, 295–308 (2011).

35 Bardella, G., Bifone, A., Gabrielli, A., Gozzi, A. & Squartini, T. Hierarchical organization of functional connectivity in the mouse brain: a complex network approach. Scientific Reports 6, 1–11 (2016).

36 Keller-Ressel, M. & Nargang, S. Hydra: a method for strain-minimizing hyperbolic embedding of network-and distance-based data. Journal of Complex Networks 8, cnaa002 (2020).

37 Muscoloni, A. & Cannistraci, C. V. A nonuniform popularity-similarity optimization (nPSO) model to efficiently generate realistic complex networks with communities. New Journal of Physics 20, 052002 (2018).

38 Papadopoulos, F., Psomas, C. & Krioukov, D. Network mapping by replaying hyperbolic growth. IEEE/ACM Transactions on Networking 23, 198–211 (2014).

39 Suzuki, R., Takahama, R. & Onoda, S. Hyperbolic disk embeddings for directed acyclic graphs. in International Conference on Machine Learning. 6066–6075 (PMLR).

40 Papadopoulos, F., Aldecoa, R. & Krioukov, D. Network geometry inference using common neighbors. Physical Review E 92, 022807 (2015).

41 Alanis-Lobato, G., Mier, P. & Andrade-Navarro, M. A. Manifold learning and maximum likelihood estimation for hyperbolic network embedding. Applied network science 1, 1–14 (2016).

42 Power, J. D. et al. Functional network organization of the human brain. Neuron 72, 665–678 (2011).

43 Mancuso, L. et al. The homotopic connectivity of the functional brain: a meta-analytic approach. Scientific Reports 9, 3346, doi:10.1038.s41598-019-40188-3 (2019).

44 Wei, P., Zhang, Z., Lv, Z. & Jing, B. Strong Functional Connectivity among Homotopic Brain Areas Is Vital for Motor Control in Unilateral Limb Movement. Frontiers in Human Neuroscience 11, doi:10.3389.fnhum.2017.00366 (2017).

45 Lee, D. S. et al. Metabolic connectivity by interregional correlation analysis using statistical parametric mapping (SPM) and FDG brain PET; methodological development and patterns of metabolic connectivity in adults. European Journal of Nuclear Medicine and Molecular Imaging 35, 1681–1691 (2008).

46 Ganea, O., Bécigneul, G. & Hofmann, T. Hyperbolic entailment cones for learning hierarchical embeddings. in International Conference on Machine Learning. 1646–1655 (PMLR).

47 Alvarez-Melis, D., Mroueh, Y. & Jaakkola, T. Unsupervised hierarchy matching with optimal transport over hyperbolic spaces. in International Conference on Artificial Intelligence and Statistics. 1606–1617 (PMLR).

48 Baio, J. et al. Prevalence of autism spectrum disorder among children aged 8 years—autism and developmental disabilities monitoring network, 11 sites, United States, 2014. MMWR Surveillance Summaries 67, 1 (2018).

49 Belmonte, M. K. et al. Autism and abnormal development of brain connectivity. Journal of Neuroscience 24, 9228–9231 (2004).

50 Bullmore, E. & Sporns, O. The economy of brain network organization. Nature Reviews Neuroscience 13, 336–349 (2012).

51 Just, M. A., Cherkassky, V. L., Keller, T. A. & Minshew, N. J. Cortical activation and synchronization during sentence comprehension in high-functioning autism: evidence of underconnectivity. Brain 127, 1811–1821 (2004).

52 Maximo, J. O., Cadena, E. J. & Kana, R. K. The implications of brain connectivity in the neuropsychology of autism. Neuropsychology Review 24, 16–31 (2014).

53 Kleinhans, N. M. et al. Abnormal functional connectivity in autism spectrum disorders during face processing. Brain 131, 1000–1012 (2008).

54 Green, S. A., Hernandez, L., Bookheimer, S. Y. & Dapretto, M. Salience network connectivity in autism is related to brain and behavioral markers of sensory overresponsivity. Journal of the American Academy of Child Adolescent Psychiatry 55, 618–626. e611 (2016).

55 Uddin, L. Q. et al. Salience network–based classification and prediction of symptom severity in children with autism. JAMA psychiatry 70, 869–879 (2013).

56 Wing, L. Asperger’s syndrome: a clinical account. Psychological Medicine 11, 115–129 (1981).

57 Frith, U. & Mira, M. Autism and Asperger syndrome. Focus on Autistic Behavior 7, 13–15 (1992).

58 Isik, L., Koldewyn, K., Beeler, D. & Kanwisher, N. Perceiving social interactions in the posterior superior temporal sulcus. Proceedings of the National Academy of Sciences 114, E9145–E9152 (2017).

59 Materna, S., Dicke, P. W. & Thier, P. Dissociable roles of the superior temporal sulcus and the intraparietal sulcus in joint attention: a functional magnetic resonance imaging study. Journal of Cognitive Neuroscience 20, 108–119 (2008).

60 Fox, M. D., Zhang, D., Snyder, A. Z. & Raichle, M. E. The global signal and observed anticorrelated resting state brain networks. Journal of Neurophysiology 101, 3270–3283 (2009).

61 Liang, Z., King, J. & Zhang, N. Anticorrelated resting-state functional connectivity in awake rat brain. Neuroimage 59, 1190–1199 (2012).

62 Van Essen, D. C. et al. The Human Connectome Project: a data acquisition perspective. Neuroimage 62, 2222–2231 (2012).

63 Di Martino, A. et al. Enhancing studies of the connectome in autism using the autism brain imaging data exchange II. Scientific Data 4, 1–15 (2017).

64 Fan, L. et al. The human brainnetome atlas: a new brain atlas based on connectional architecture. Cerebral Cortex 26, 3508–3526 (2016).

65 Cui, P., Wang, X., Pei, J. & Zhu, W. A survey on network embedding. IEEE Transactions on Knowledge Data Engineering 31, 833–852 (2018).

66 Nickel, M. & Kiela, D. Poincaré embeddings for learning hierarchical representations. arXiv preprint arXiv:.08039 (2017).

67 García-Pérez, G., Aliakbarisani, R., Ghasemi, A. & Serrano, M. Á. Precision as a measure of predictability of missing links in real networks. Physical Review E 101, 052318 (2020).

68 van de Ven, V. G., Formisano, E., Prvulovic, D., Roeder, C. H. & Linden, D. E. Functional connectivity as revealed by spatial independent component analysis of fMRI measurements during rest. Human Brain Mapping 22, 165–178 (2004).

