## Supplementary Information for "Hyperbolic disc embedding of functional human brain connectomes using resting state fMRI"

**Contents**

**Supplementary Fig. 1** Voxel-wise embedding onto hyperbolic discs by $\mathbb{S}^{1}$/$\mathbb{H}^{2}$ model

**Supplementary Fig. 2** Plotting of independent components (ICs) using colors upon the voxel-wise hyperbolic embedded discs

**Supplementary Fig. 3** Comparison of embedded hyperbolic discs of voxel correlational structures per each independent component (IC) over 10 subjects

**Supplementary Fig. 4** Mean, standard deviation (SD), coefficient of variation (CV) of hyperbolic distance analog of a representative case, which were derived from 100 times-repeated hyperbolic embedding on $\mathbb{S}^{1}$/$\mathbb{H}^{2}$ model

**Supplementary Note 1** Comparison of pattern of distribution between absolute correlation (derived from time-series correlation) and connection probability (after hyperbolic embedding and distance calculation between node pairs)

**Supplementary Note 2** Coefficient of variation (CV) of repeated hyperbolic embedding and the hyperbolic distance analog between node pairs on this embedded hyperbolic disc

**Supplementary Note 3** Brief description of the geometric 𝕊^1^/ ℍ^2^ model derived from Serrano et al.^1^ and Krioukov et al.^2^

**Supplementary Note 4** List of abbreviations in Fig. 7

**References**

**Supplementary Figures**

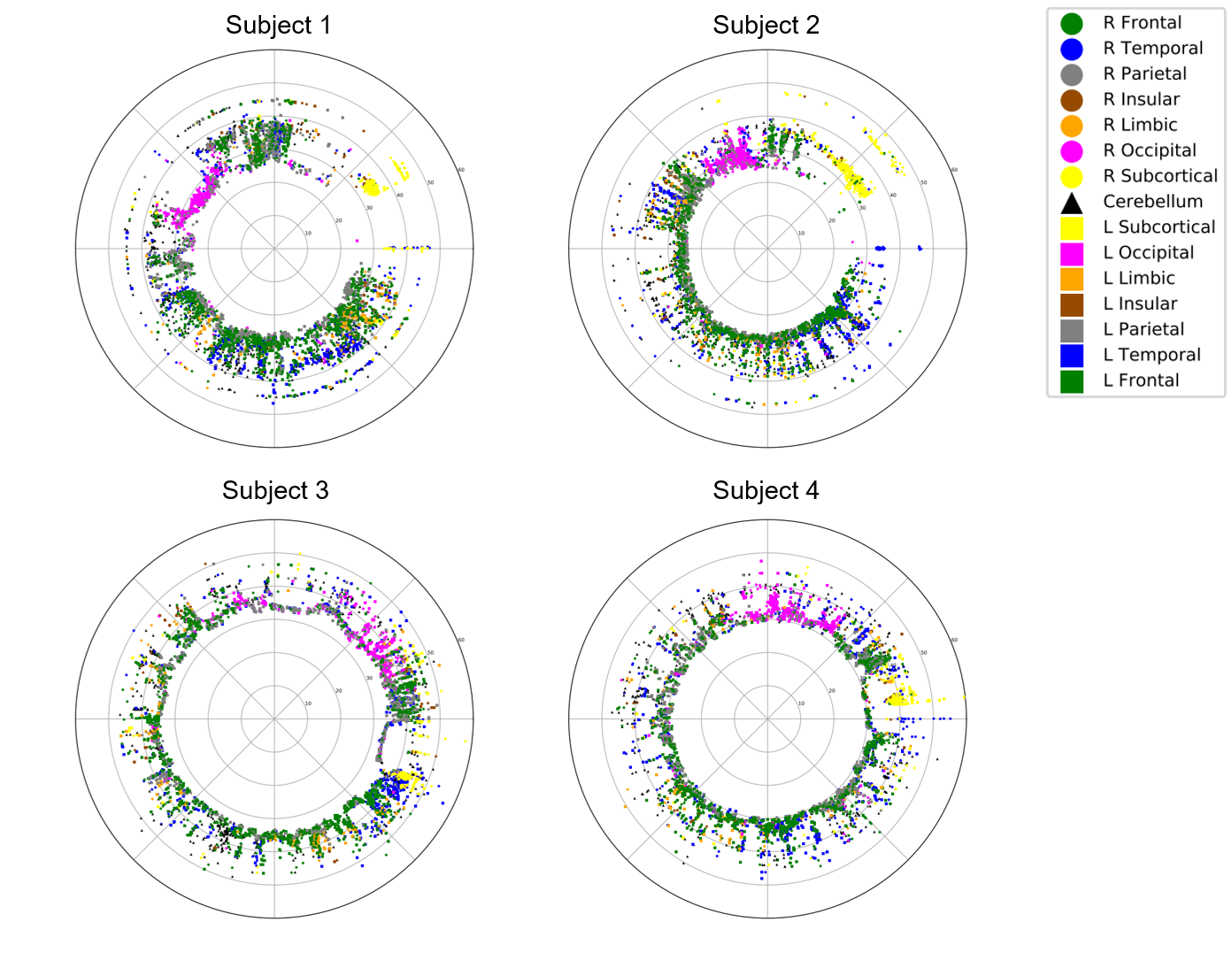

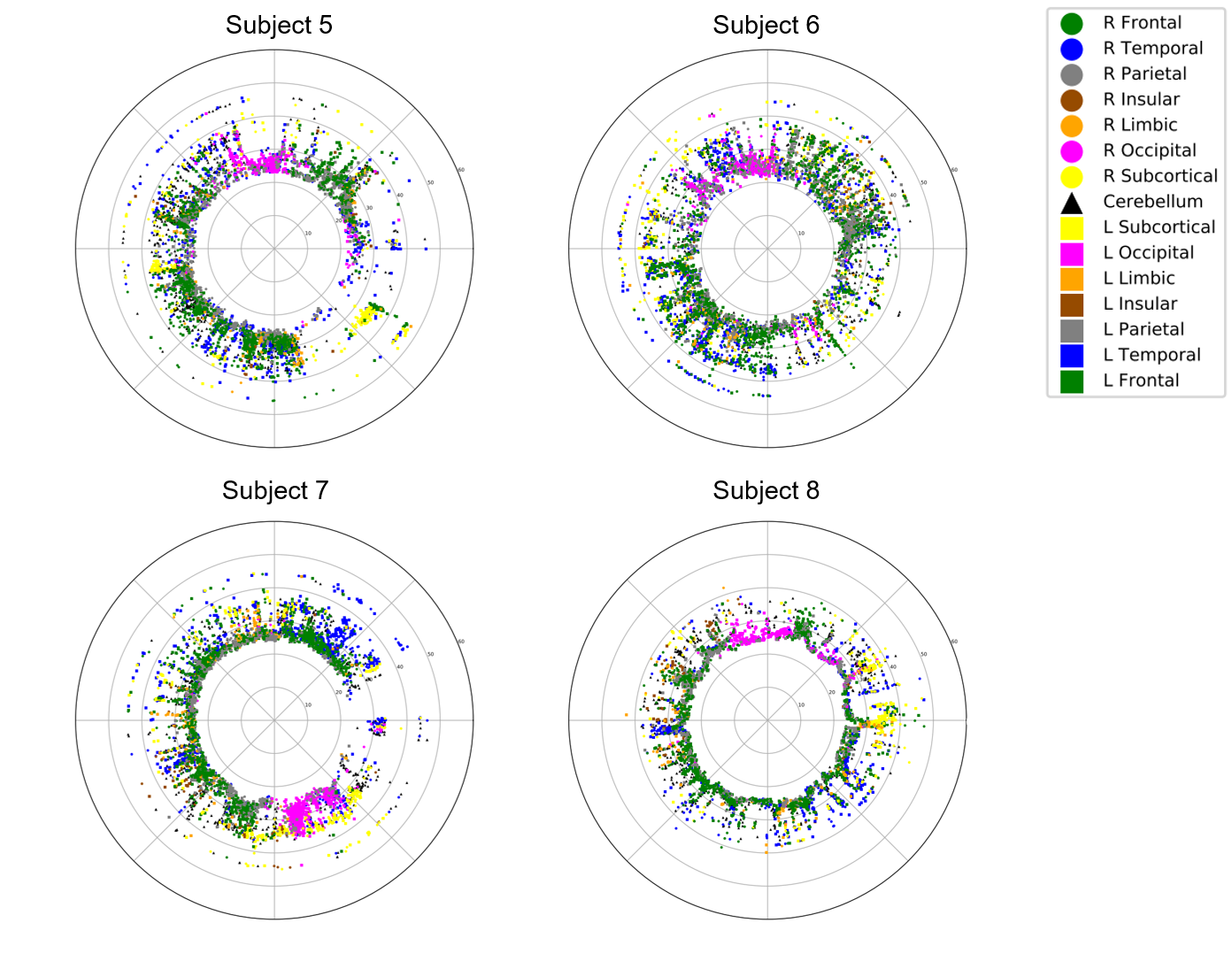

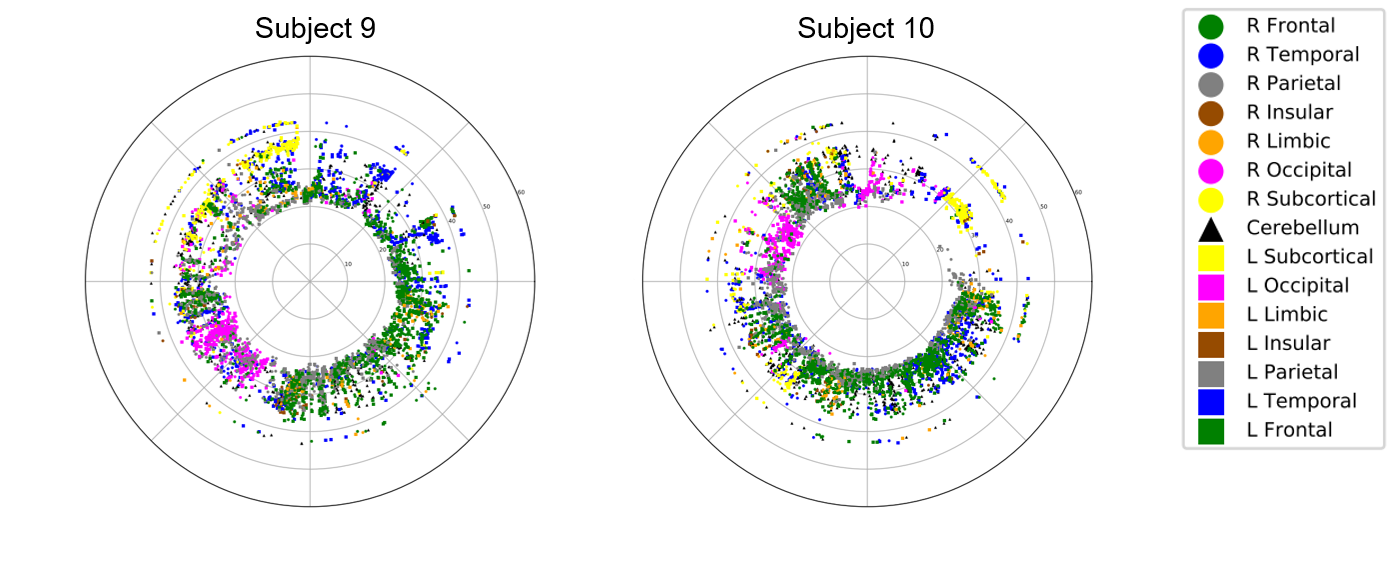

**Supplementary Fig. 1 Voxel-wise embedding onto hyperbolic discs by** $\mathbb{S}^{1}$/$\mathbb{H}^{2}$ **model.** Color denotes the anatomic lobe in which each region of interest (ROI) is located, while the lobes within the right, left cerebral hemisphere, and cerebellum are marked with circle, square and triangular markers, respectively. Similar to the result in ROI-scope, vacant space in the center of disc was prominent. Nodes in the same anatomic lobe tended to distribute in similar angular coordinate.

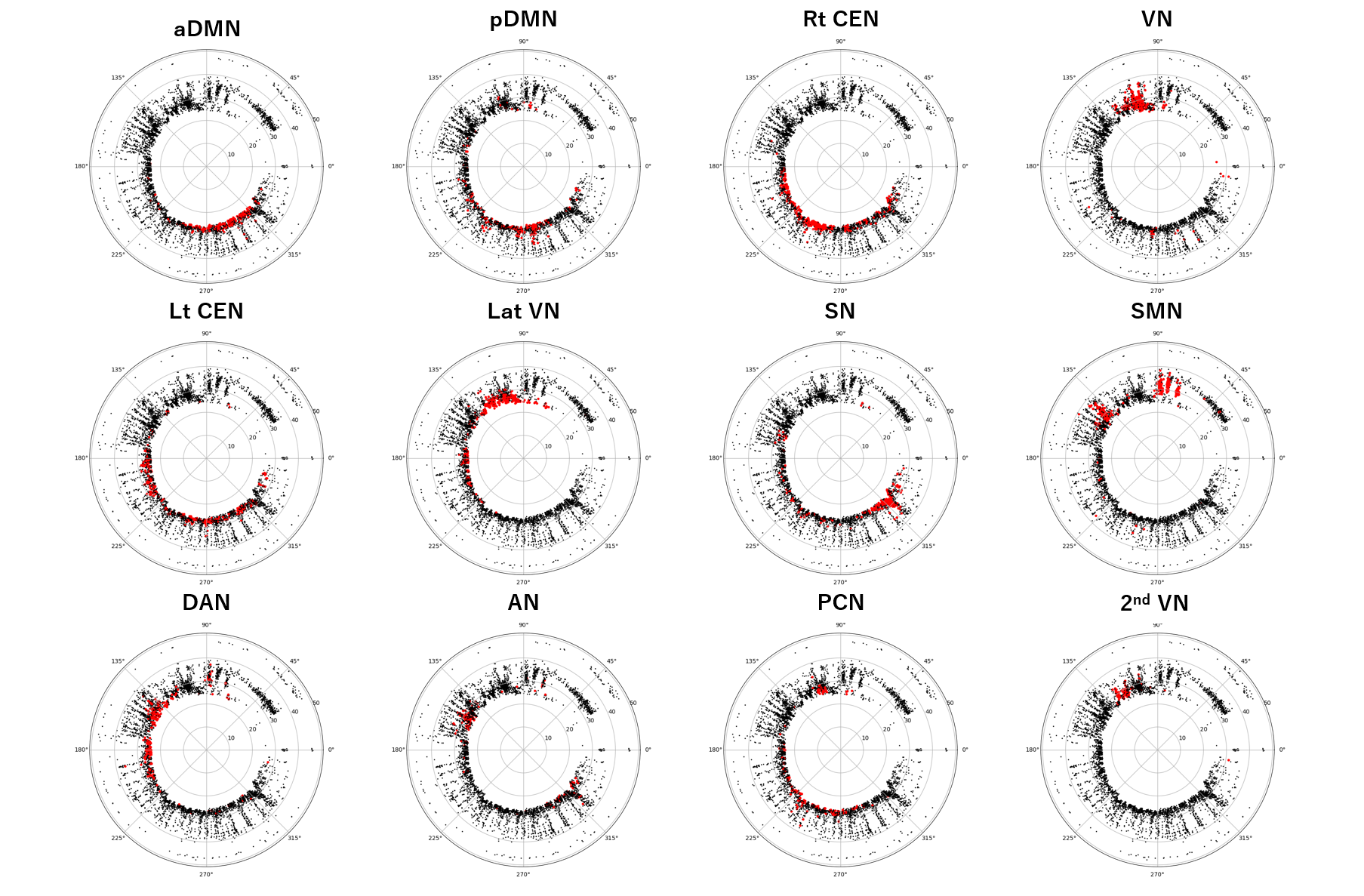

Supplementary Fig. 2 Plotting of independent components (ICs) using colors upon the voxel-wise hyperbolic embedded discs. Using voxel data of HCP database, group-wise independent component analysis (ICA) was performed to yield 12 independent components (ICs). Voxel-wise embedded hyperbolic discs from 10 individual subjects were plotted with these ICs in color on the hyperbolic discs. A single representative case was used for visualization for this Figure. Voxels colored in red indicate the voxels included in each independent component (IC). aDMN = anterior default mode network, pDMN = posterior default mode network, CEN = central executive network, VN = visual network, Lat VN = lateral visual network, 2^nd^ VN = secondary visual network, SN = salience network, SMN = sensorimotor network, DAN = dorsal attention network, AN = auditory network, PCN = precuneus network.

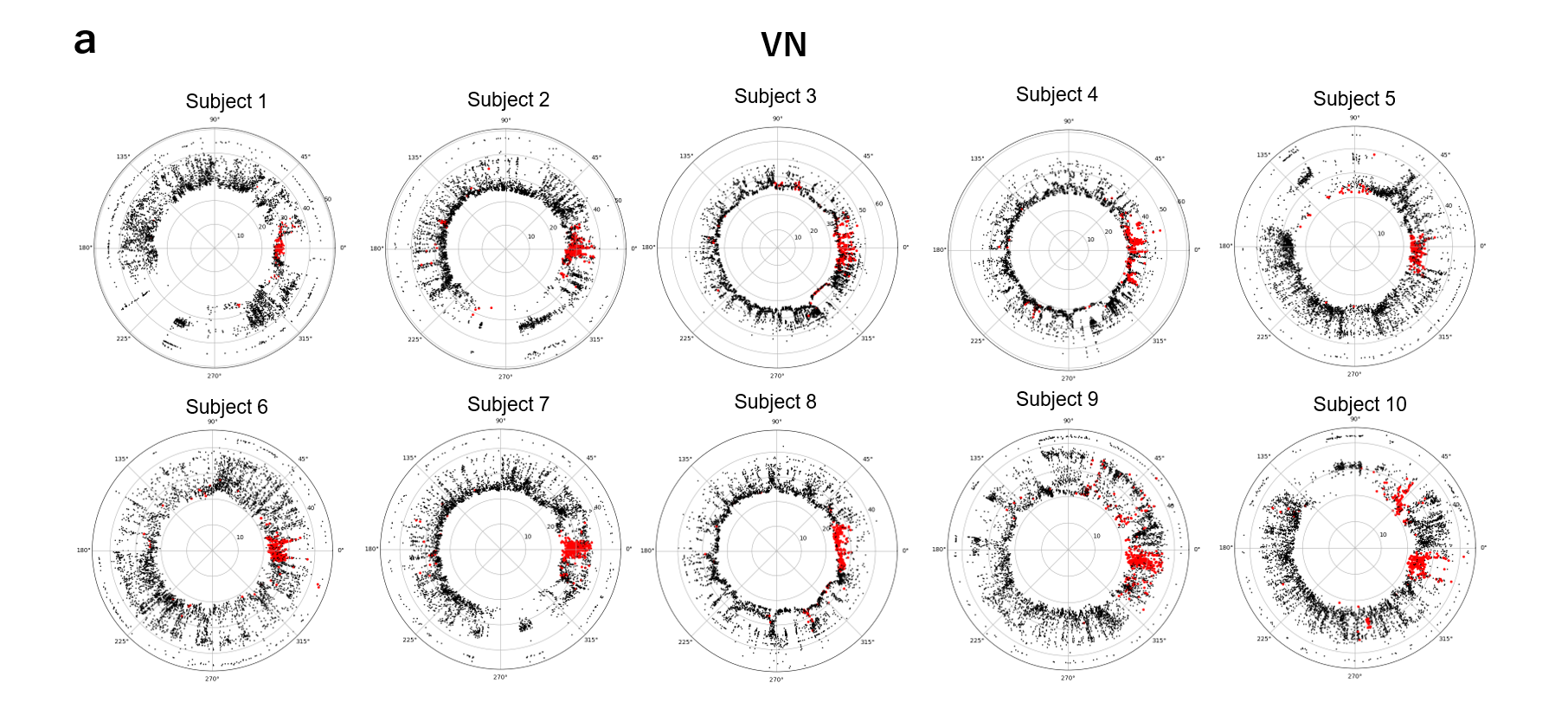

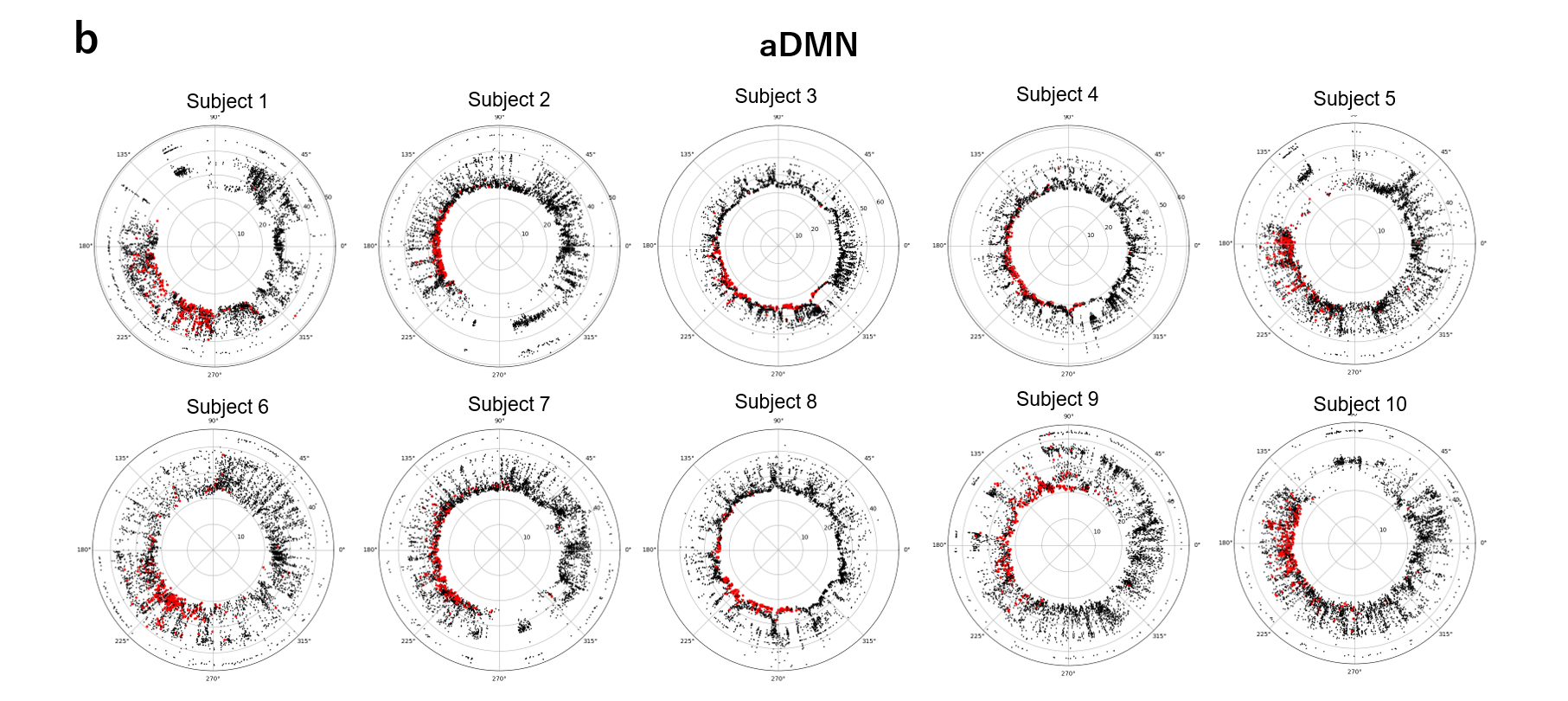

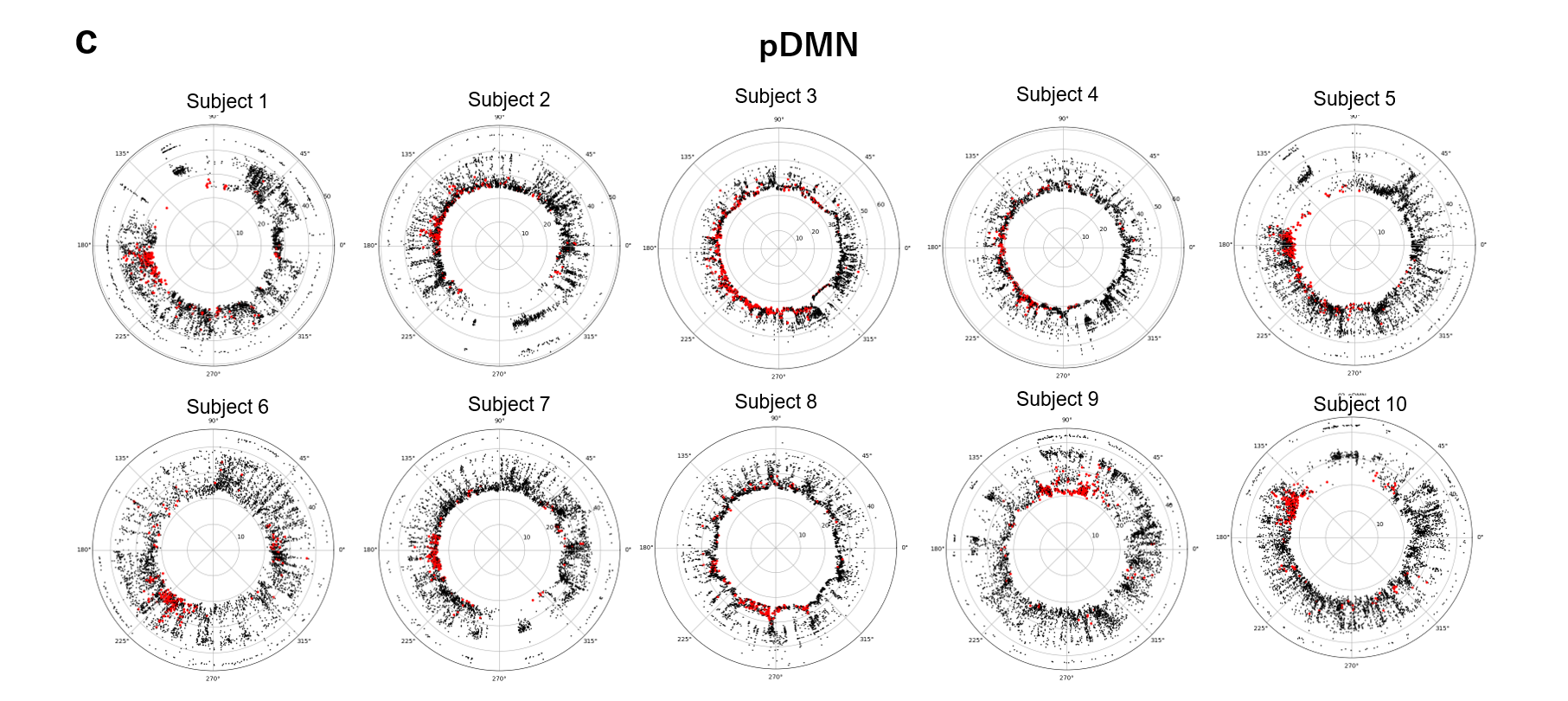

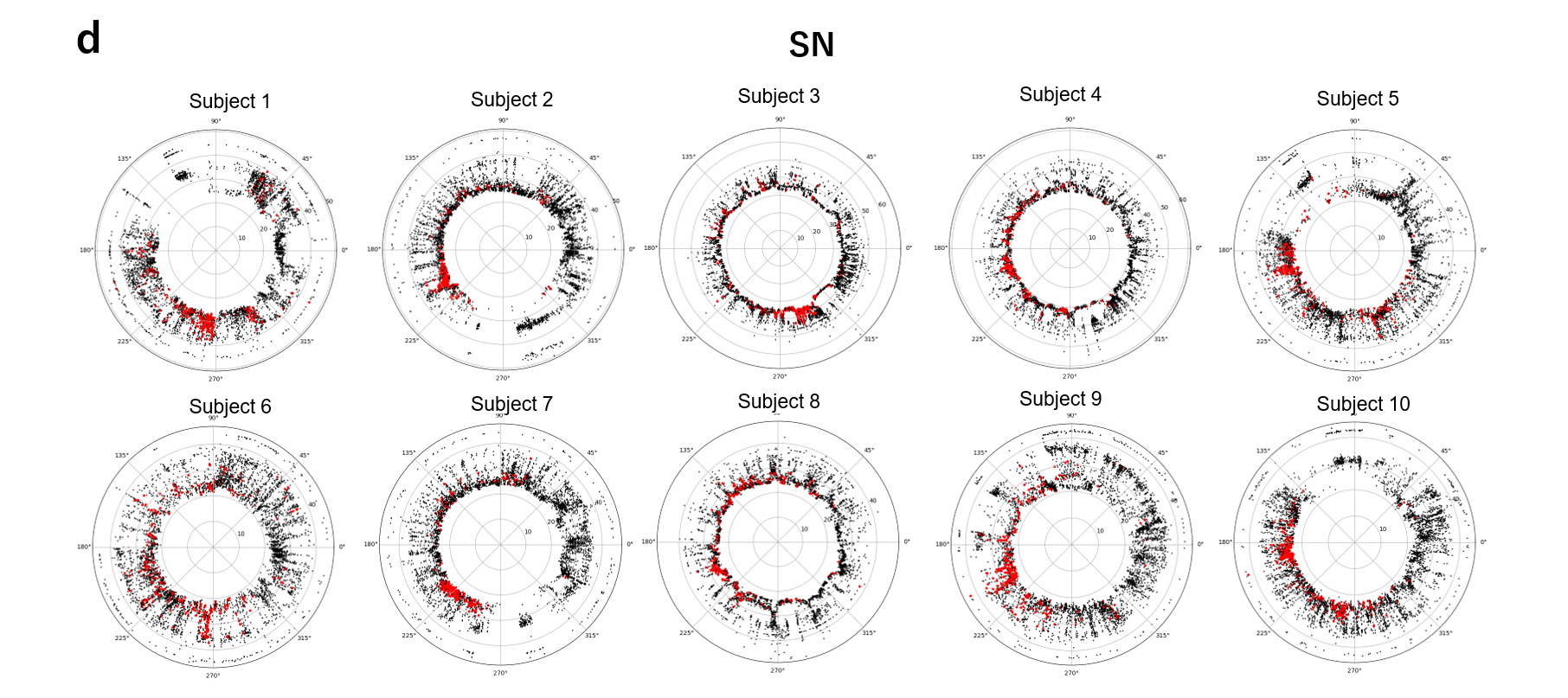

**Supplementary Fig. 3 Comparison of embedded hyperbolic discs of voxel correlational structures per each independent components (ICs) over 10 subjects.** Absolute angular position of the mean angle of visual network (VN) was set arbitrarily to have a value of angle 0 (right direction of the figure) on the embedded hyperbolic disc. Thus, VN was located in the same angle, however, the angular coherence seemed to be higher for this IC VN. Colored independent components such as anterior DMN and posterior DMN in each subject showed ‘grossly similar but different in detail’ pattern of wider distribution than VN along individuals. IC of SN showed a broader distribution along the 10 individuals. IC: independent component, VN: visual network, DMN: default mode network, SN: salience network.

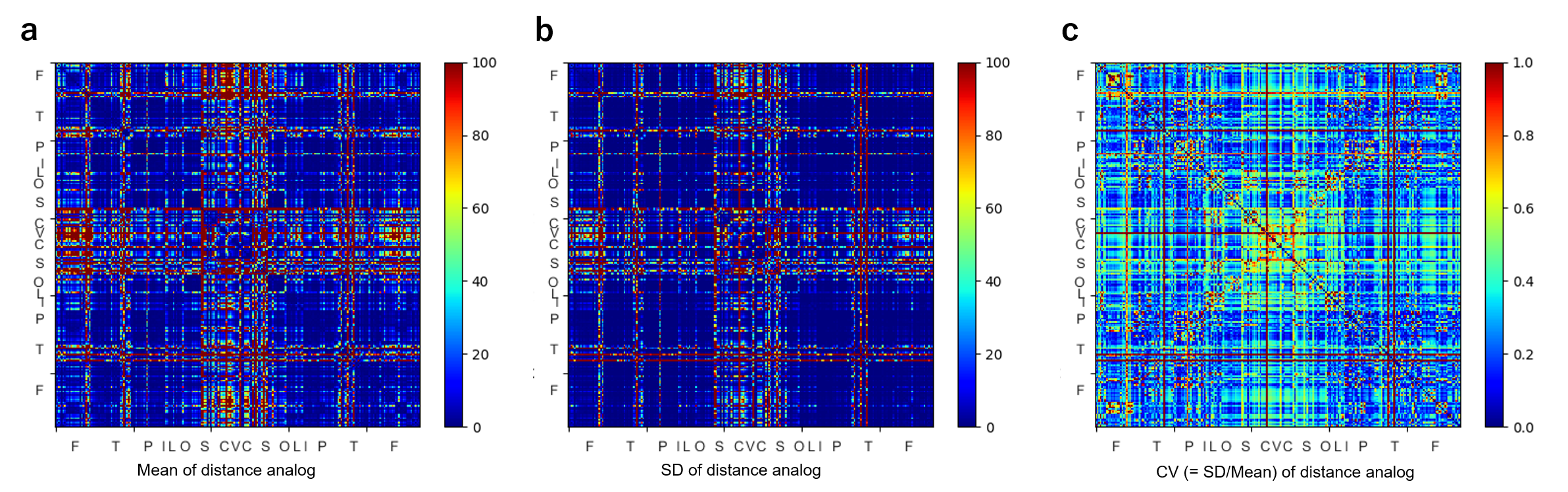

**Supplementary Fig. 4 Mean, standard deviation (SD), coefficient of variation (CV) of hyperbolic distance analog of a representative case, which were derived from 100 times-repeated hyperbolic embedding on** $\mathbb{S}^{\boldsymbol{1}}$**/**$\mathbb{H}^{\boldsymbol{2}}$ **model.** As a descriptive measure of how the hyperbolic length of an edge on embedded hyperbolic disc varies over multiple times of embedding procedure, we used the coefficient of variation (CV) of the hyperbolic distance analog. **a** Mean, **b** standard deviation (SD), **c** coefficient of variation (CV) of distance analog over repetition of embedding is represented in the figure. (Subfigure **c** is same with **Fig. 6b** in the main text but duplicated for the sake of comparability.) Edges with longer distance tended to have bigger variance over repetition of embedding, and some of them had also larger CV value (**Supplementary Note 2**).

**Supplementary Notes**

1. **Comparison of pattern of distribution between absolute correlation (derived from time-series correlation) and connection probability (after hyperbolic embedding and distance calculation between node pairs)**

In this work, we made use of two proximity measures between two nodes. First is the absolute correlation coefficient, which we used to compose the adjacency matrix. Second is the connection probability, which was computed from the hyperbolic distance that resulted from the hyperbolic disc embedding.

Note that the two measures are ranged from 0 to 1, and both get closer to 1 as the two nodes are closely correlated (or functionally similar). However, since the correlation is computed from the quadratic formula of deviation as in Eq. (3) and the connection probability is calculated from the exponential of hyperbolic distance, it is difficult to compare the proximity computed by two measures analytically.

Nevertheless, we can address the similarity of *pattern* of distribution of two proximity measures. To quantitatively assess how similar the two proximity measures are distributed, we computed the root mean square of deviation $S$, given as the following:

$$S= \sqrt{\frac{\sum_{i>j} \left| p_{ij}-\left| \hat{\rho}_{ij} \right| \right|^{2}}{N\left( N-1 \right)/2}-\left| \bar{p}-\bar{\left| \hat{\rho} \right|} \right|^{2}}$$

(S1)

where $\bar{p}$ and $\bar{|\hat{\rho}|}$ denotes the mean value of connection probability and absolute correlation over all the edges. $N$ is the number of nodes included in the component, $p_{ij}$ and $\hat{\rho}_{ij}$ denotes connection probability and Pearson’s correlation for edges connecting node $i$ and $j$, respectively. The first term in the square root is the mean square of the edge-wise difference between two measures, while the second term is the difference between mean values of connection probability and absolute correlation value.

Therefore, by Eq. (S1), we can assess the similarity of pattern between two measures, by observing how much larger the mean square of the edge-wise difference between two measures would be than that indicated by the mean difference between two matrices. As $S$ gets smaller, two distributions can be said more similar. Note that $S$ is no less then zero, as easily shown by elementary statistics. The equality of the first and the second term (and thus deviation S = 0) is satisfied when the differences of elements are of constant values over all the edges.

Values of $S$ among 180 subjects from HCP dataset were distributed as Mean = 0.21, SD = 0.062 among subject. For the subjects shown in the (**Fig. 5**) of the main text, the value is 0.18. 0.14, 0.15, and 0.13 for subject 1, 2, 3, and 4, respectively.

When we computed the same measure with the connection probabilities inter-individually among 180 subjects (total of $(180 \times179) / 2$ pairs), the values of $S$ were distributed as Mean = 0.29, SD = 0.035 among all pairs of subjects. Compared to this inter-individual difference of deviation S values of connection probability between subject pairs, difference of the two proximity measures within individuals had significantly more similar pattern (*P* < 0.001). In this manner, we could address the two proximity measures had the similarity, though not the same, as described in the main text.

2. **Coefficient of variation (CV) of repeated hyperbolic embedding and the hyperbolic distance analog between node pairs on this embedded hyperbolic disc**

As a descriptive measure of how the hyperbolic length of an edge varies over multiple times of embedding procedure, we used the coefficient of variation (CV), which is a well-known measure of evaluating dispersion, computed by ratio of standard deviation to mean, of the distance analog computed by Eq. (8) in the main text.

$$D_{ij}=e^{\frac{\left( d_{ij}-\hat{R} \right)}{2}}$$

Though the distance analog is not a linear parameter, the CV measure suffices to compare between edges to evaluate which edge has more aberrant distribution over multiple instances of embedding.

For the representative case same as in (**Fig. 6**) in the main text, we demonstrated the mean, standard deviation (SD) and CV values in (**Supplementary Fig. 4**). Each element of matrix denotes edge. Therefore, the horizontal or vertical line denotes the edges connected to the same node. The red-colored lines shown in the (**Supplementary Fig. 4a**) suggests that all of the edges that are connected to those specific nodes with red elements (edges) have larger hyperbolic distance than others with green/blue/black edges, which means that those red-lined nodes are located farther from most of the other nodes with green/blue/black lines. The edges connected to these nodes also have high standard deviation (**Supplementary Fig. 4b**). And some, but not all, of these lines reappear at the (**Supplementary Fig. 4c**) which suggests higher dispersion meaning non-reproducibility along embedding.

Therefore, we could address, from this case presentation, that some of the nodes that are located on the embedded hyperbolic discs farther from most of the other nodes, which also indicates their non-popularity (very low degree) in functional brain network, have higher arbitrariness over the repetition of embedding.

3. **Brief description of the geometric 𝕊^1^/ ℍ^2^ model derived from Serrano et al.^1^ and Krioukov et al.^2^**
A. The similarity space: 𝕊^1^ model

The similarity space 𝕊^1^ is a one-dimensional sphere^1^, which is equivalent to a circle of radius $R$. Total number of $N$ nodes are distributed in this circle with density of 1 without loss of generality, so that the radius of circle $R=\frac{N}{2\pi}$.

$N$ nodes are also assigned a hidden variable, $\kappa$, which is proportional to its expected degree. We can assume that the $\kappa$ and angular position $\theta$ are correlated and distributed according to a distribution function of $\rho\left( \kappa,\theta\right)$. This kind of model is able to generate community structures and clustering.

If all $N$ nodes are assigned variable $\left( \kappa_{1},\theta_{1} \right),\left( \kappa_{2},\theta_{2} \right),\cdots\left( \kappa_{N},\theta_{N} \right)$, the model propose that each pair of nodes $i, j$ is connected, with probability

$$p_{ij}=\frac{1}{1+\left( \frac{d_{ij}}{\mu\kappa_{i}\kappa_{j}} \right)^{\beta}}$$

(S2)

where $d_{ij}=R\Delta\theta_{ij}$ is the arc length between two nodes $i$ and $j$, with angular distance $\Delta\theta_{ij}$. Two global parameters, $\mu$ and $\beta$ is associated with average degree and global clustering coefficient (or inverse temperature analog), respectively.

Connection probability can be any integrable function of $\chi=\frac{d}{\mu\kappa_{i}\kappa_{j}}$^1^, but in this model we implement the Fermi-Dirac distribution model, which defines geometric graphs that are clustered, small-world and with heterogeneous degree distribution^2,3^.

If we assume that nodes are uniformly distributed along the 𝕊^1^ space, we obtain the distribution $\rho\left( \kappa,\theta\right)=\frac{\rho\left( \kappa\right)}{2\pi}$ and if we set $\mu=\frac{\beta}{2\pi\left\langle k \right\rangle}\sin\frac{\pi}{\beta}$, we obtain expected degree of a node $\bar{k}\left( \kappa\right)=\kappa$ and the average degree of network $\langle k\rangle= \langle\kappa\rangle$. This justifies the association of hidden variable $\kappa$ with the expected degree.

As a result, the process of embedding is equivalent to estimating the $2N+1$ parameters, $\left( \kappa_{1},\theta_{1} \right),\left( \kappa_{2},\theta_{2} \right),\cdots\left( \kappa_{N},\theta_{N} \right)$ and $\beta$.

B. Pure geometric model: ℍ^2^ model

The 𝕊^1^ model can translated into a pure geometric model in the hyperbolic plane, by converting the expected degree $\kappa$ to the radial coordinate $r$, by the following:

$$r_{i}=\hat{R}-2\ln\frac{\kappa_{i}}{\kappa_{0}}$$

(S3)

where $\hat{R}\equiv2\ln\frac{N}{\mu\pi\kappa_{0}^{2}}.$Then, the connection probability noted in Eq. (S2) is equivalent to:

$$p_{ij}=\frac{1}{1+e^{\frac{\beta}{2}\left( d_{ij}-\hat{R} \right)}}$$

(S4)

where

$$d_{ij}\cong r_{i}+r_{j}+2\ln\frac{\Delta\theta_{ij}}{2} +O\left( \frac{1}{r} \right)$$

(S5)

is a good approximation of the distance between two nodes $i$ and $j$ in the hyperbolic space^[2]^. The hyperbolic law of cosine was used when $\Delta\theta_{ij}$ is very small.

Thus, the connection probability $p_{ij}$ becomes a single-variable function of hyperbolic distance $d_{ij}$, which makes the model purely geometric. This has important consequences for the global connectivity of network, for instance, the geodesic curves in the hyperbolic plane can be used to efficiently navigate the network ^4^.

4. **List of abbreviations in Fig. 7**

Suffix _R, _L denotes right and left hemisphere.

**Frontal lobe**

SFG = Superior Frontal Gyrus

MFG = Middle Frontal Gyrus

IFG = Inferior Frontal Gyrus
ORG = Orbital Gyrus

PrG = Precentral Gyrus

PCL = Paracentral Lobule

**Temporal Lobe**

STG = Superior Temporal Gyrus

MTG = Middle Temporal Gyrus

ITG = Inferior Temporal Gyrus

FuG = Fusiform Gyrus

PhG = Parahippocampal Gyrus

pSTS = Posterior Superior Temporal Sulcus

**Parietal Lobe**

SPL = Superior Parietal Lobule

IPL = Inferior Parietal Lobule

PCun = Precuneus

PoG = Postcentral Gyrus

**Insula Lobe**

INS = Insula

**Limbic lobe**

CG = Cingulate Gyrus

**Occipital Lobe**

MVOcC = MedioVentral Occipital Cortex

LOcC = Lateral Occipital Cortex

**Subcortical Nuclei**

Amyg = Amygdala

Hipp = Hippocampus

BG = Basal Ganglia

Tha = Thalamus

**Cerebellum and Vermis**

CB = Cerebellum

Vermis

**References**

1	Serrano, M. A., Krioukov, D. & Boguná, M. Self-similarity of complex networks and hidden metric spaces. *Physical review letters* **100**, 078701 (2008).

2. Krioukov, D., Papadopoulos, F., Kitsak, M., Vahdat, A., & Boguná, M. (2010). Hyperbolic geometry of complex networks. *Physical Review E*, **82**, 036106.

3	García-Pérez, G., Allard, A., Serrano, M. Á. & Boguñá, M. Mercator: uncovering faithful hyperbolic embeddings of complex networks. *New Journal of Physics* **21**, 123033 (2019).

4	Boguna, M., Krioukov, D. & Claffy, K. C. Navigability of complex networks. *Nature Physics* **5**, 74-80 (2009).

2. ^a^ The approximation is accurate for two nodes that are separated by $\Delta\theta_{ij}>\sqrt{e^{-2r_{i}}+e^{-2r_{j}}}$
